# Genetic dissection of femoral and tibial microarchitecture

**DOI:** 10.1101/512103

**Authors:** Lu Lu, Jinsong Huang, Fuyi Xu, Zhousheng Xiao, Jing Wang, Bing Zhang, Nicolae Valentin David, Danny Arends, Weikuan Gu, Cheryl Ackert-Bicknell, Olivia L. Sabik, Charles R. Farber, Leigh Darryl Quarles, Robert W. Williams

**Affiliations:** Department of Genetics, Genomics and Informatics, University of Tennessee Health Science Center, Memphis, TN, USA; Department of Medicine, University of Tennessee Health Science Center, Memphis, TN, USA; Department of Molecular and Human Genetics, Baylor College of Medicine, Houston, TX, USA; Department of Medicine, Northwestern University Feinberg School of Medicine, Chicago, IL, USA; Züchtungsbiologie und molekulare Tierzüchtung, Humboldt University, Berlin, Germany; Department of Orthopaedic Surgery and Biomedical Engineering, University of Tennessee Health Science Center, Memphis, TN, USA; Center for Musculoskeletal Research, University of Rochester, Rochester, NY, USA; Center for Public Health Genomics, University of Virginia, Charlottesville, VA, USA

**Keywords:** microCT, trabecular bone, cortical bone, QTL, gene ontology, systems genetics, animal model, GWAS, ignorome

## Abstract

Our understanding of the genetic control of bone has relied almost exclusively on estimates of bone mineral density. In contrast, here we have used high-resolution x-ray tomography (8 μm isotropic voxels) to measure femoral and tibial components across a set of ~600 mice belonging to 60 diverse BXD strains of mice. We computed heritabilities of 25 cortical and trabecular compartments. Males and females have well matched trait heritabilities, ranging from 0.25 to 0.75. We mapped 16 QTLs that collectively cover ~8% of all protein-coding genes in mouse. A majority of loci are detected only in females, and there is also a bias in favor of QTLs for cortical traits. To efficiently evaluate candidate genes we developed a method that couples gene ontologies with expression data to compute bone-enrichment scores for almost all protein-coding genes. We carefully collated and aligned murine candidates with recent human BMD genome-wide association results. We highlight a subset of 50 strong candidates that fall into three categories: 1. those linked to bone function that have already been experimentally validated (*Adamts4, Ddr2, Darc, Adam12, Fkbp10, E2f6, Adam17, Grem2, Ifi204*); 2. candidates with putative bone function but not yet tested (e.g., *Greb1, Ifi202b*) but several of which have been linked to phenotypes in humans; and 3. candidates that have high bone-enrichment scores but for which there is not yet any specific link to bone biology or skeletal disease, including *Ifi202b, Ly9, Ifi205, Mgmt, F2rl1, Iqgap2*. Our results highlight contrasting genetic architecture between the sexes and among major bone compartments. The joint use and alignment of murine and human data should greatly facilitate function analysis and preclinical testing.

**Disclosure:** The authors declare that no competing interests exist.

## Introduction

The development and maintenance of the skeletal system is modulated by a large number of genetic and environmental factors with many adaptive and age-related changes in bone phenotype that lead to osteoporosis and increased fracture risk. Over the past decade more than 1000 loci and gene variants have been defined using human, mouse, and rat cohorts that control bone mineral density, risk of fracture, and other morphometric traits ^(1)^. Most large genetic studies of osteoporosis in humans have exploited dual-energy x-ray absorptiometry (DXA) to quantify areal bone mineral density (BMD) ^(2)^. Some of genome-wide association studies (GWAS) have exploited peripheral quantitative computed tomography (pQCT) to define subsets of sequence variants that modulate bone architecture and strength ^(3)^. While BMD accounts for about 70% of bone strength, this method lacks the 3-dimensional (3D) structural precision of high-resolution microCT ^(4–7)^. 3D maps of structure and architecture generated by microCT have many advantages, including (1) avoidance of interference from intra- and extra-osseous soft tissues, (2) high-content data acquisition, and (3) isotropic resolution as high as 6 μm ^(8, 9)^. Finally, the development of finite element analysis models derived from microCT data provide a way to model mechanical properties of bone ^(10)^.

Experimental rodent models provide a way to evaluate candidate genes generated in GWAS and other human genetic and genomic studies and to reduce variants to mechanisms and even treatments ^(11)^. The use of knockouts or knockins of single mutation is a well-established approach to test the roles of genes on skeletal system structure and function. An alternative unbiased whole genome approach to map natural genetic variants that control bone growth and homeostasis uses a large family panel of genetically diverse mice ^(12)^. This allows some of the clinical complexity of bone disease to be captured, while retaining tight control over diet, environment, and genotypes. For example, the set of 96 strains that are part of the Hybrid Mouse Diversity Panel ^(13, 14)^ and a subset of strains that belong to the BXD recombinant inbred (RI) family have been used to define a key role of *ASXL2* in BMD and an important role of *ALPL* in hypophosphatasia ^(15)^. In comparison to standard F2 intercrosses, these large families of isogenic but diverse strains can be used to systematically test gene-by-environmental interactions, to evaluate the replicability of findings, and to test new therapies and treatments ^(16)^. Families of strains, such as the BXDs, are also advantageous because so much genomic, metabolic, and phenotypic data has already been collected for these mice ^(17)^. It becomes practical to measure heritability for virtually any trait and to map sets of quantitative trait loci (QTLs) that influence bone and other traits ^(15, 18–21)^.

In this study we have combined deep quantitative phenotyping of bone microstructure derived from microCT combined with a fine-grained genetic dissection to understand the complex control of regional bone traits-focusing on femur and tibia. We have systematically evaluate the whole bone, and cortical and trabecular segments in the BXD family of strains. Finally, we have applied a new method to rank essentially all protein-coding genes in mice, and therefore humans, with respect to their potential roles in bone biology. We define for the first time those genes with no published function or literature connection to skeletal homeostasis as “bone ignorome” ^(22, 23)^. These set of uncharacterized genes are likely to be important in skeletal system biology and function. We have merged information on microCT-associated genes with data on the bone ignorome to systematically rank candidate that may contribute to variation in bone size and architecture.

## Materials and Methods

### Animals

All the mouse experimental procedures were in accordance with the *Guidelines for the Care and Use of Laboratory Animals* published by the National Institutes of Health and were approved by institutional Animal Care and Use Committee in the University of Tennessee Health Science Center (UTHSC).

The BXD family of strains was housed in a single specific pathogen-free (SPF) facility at UTHSC, and maintained at 20 °C to 24 °C on a 14/10 hours light/dark cycle. The mice were provided with 5% fat Agway Prolab 3000 (Agway Inc., Syracuse, NY) mouse chow and Memphis aquifer tap water, ad libitum. Sixty-one BXD strains and both parental strains, C57BL/6J (B6) and DBA/2J (D2), were sacrificed for tissue harvest. The age of these mice ranged from 50 to 375 days with an average of 100 days. A total of 597 animals were studied, including 290 females and 307 males. Five hundred and seventy-six femurs and 515 tibias were harvested. The differences in animal numbers and bone numbers reflect breakage or loss of material prior to measurement. The precise numbers of BXD strains varies by trait, sex, and age, but all parameters were evaluated using between 50 to 63 strains. (For details on sample sizes see **Supplemental Data S1** and data in GeneNetwork.org). Many cases used here had been previously used for other analysis by Zhang and colleagues ^(24)^, although here we have integrated much additional data, including 239 new cases and 15 additional strains. After dissection and removal of soft tissue, femurs and tibias were stored in 75% ethanol until measurement.

### MicroCT measurements

High-resolution x-ray tomography (μCT40, Scanco Medical, Basserdorf, Switzerland) was used to scan and measure morphometric parameters of femurs and tibias. Bones were placed in a 12.3-mm-diameter sample holder filled with 75% ethanol and immobilized with styrofoam. Samples were scanned at 8-μm resolution (isotropic voxel size) using an energy level of 55 kVp, an integration time of 300 ms, and an intensity of 109 μA. Morphometric parameters were evaluated using a fixed Gaussian filter and a threshold of 220 for trabecular bone, and 250 for both cortical bone and whole bone.

Each femur and tibia was measured separately using Scanco software that estimates values of more than 50 quantitative traits per bone (whole bone, cortical and trabecular segments). We selected a subset of 25 of the more interesting and interpretable traits for in-depth phenotyping as listed in Table 1. Three whole-bone parameters were measured: length, mineralized volume, and material bone mineral density (mBMD). For cortical bone, 100 transverse slices were acquired at the middle of the shaft—a total length of 0.8 mm. From these cross-sections a total of 11 cortical microtraits were generated, including: cortical thickness, cortical volume, porosity, polar and area moment of inertia. For trabecular bone analysis, 100 slices were acquired at the secondary spongiosa of the distal femur or the proximal tibia. A total of 11 trabecular microtraits were generated, including among others, bone volume fraction (BV/TV), trabecular thickness, trabecular number, trabecular separation, and the trabecular connectivity density (Figure 1A)

**Figure 1.**
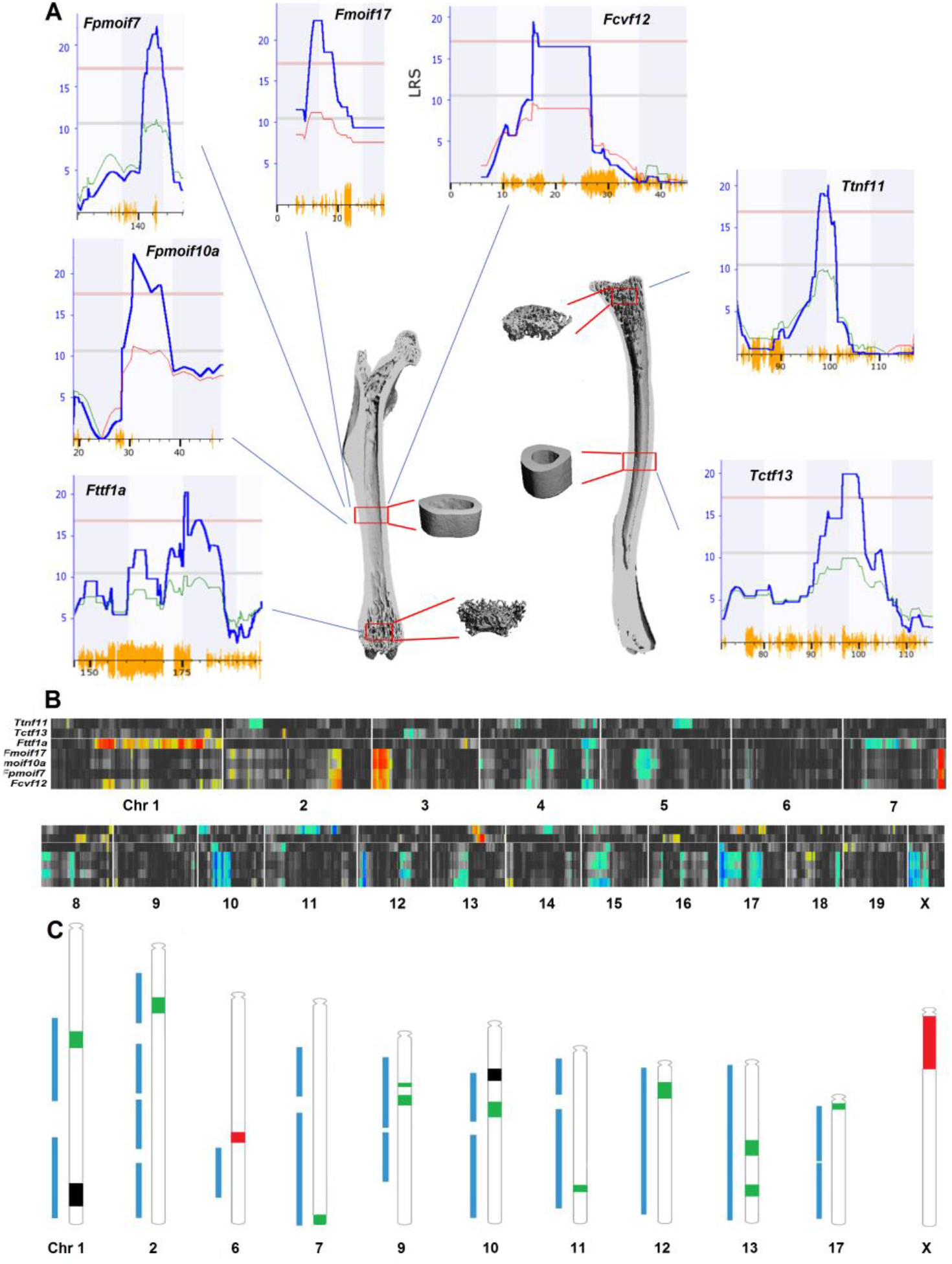
(A) Representative microCT image reconstructions of whole bone (cut-planes of femur on the left and tibia on the right). Four red boxes represent reconstructed microtraits of either cortical bone (midshaft) or trabecular bone (bottom and top, distal femur and proximal tibia) generated from 100 transverse sectional slices. The seven most robust QTLs are shown around the periphery with corresponding trait identifiers and QTLs: first letter *F* or *T* (femur or tibia), followed by abbreviation of key bone phenotype, and *f* or *m* (female or male if a QTL was sex-specific, *pmoi* = polar moment of inertia, *cv* = cortical volume, *ct* = cortical thickness, *tn* = trabecular number, *tt* = trabecular thickness). The final number is the chromosome number. For each QTL map, the x axis is given in megabases, the left y axis is the LRS score. The red horizontal line provides the genome-wide significance level based on 2,000 permutations. Orange hash along the x axis indicates SNP density. Heavy blue lines provide linkage statistics, whereas thin green and red lines provide an estimate of the additive genetic effect (right y axis). ^(130)^ for details on replicating these QTL maps. (B) The QTL heat map provides whole-genome mapping results for all seven phenotypes in the form of color-coded horizontal bands. Bands of more intense color correspond to QTL linkage peaks, and colors encode the additive effect of alleles (blue for *B* and red for *D* alleles). (C) Chromosomal ideograms for all 16 significant QTLs (Chrs 1, 2, 6, 7, 9, 10, 11, 12, 13, 17, and X) combined with mouse bone QTLs (interval coverage in blue bars) on these chromosomes listed on RGD from GViewer (www.rgd.mcw.edu/rgdweb/search/qtls.html?term=bone&chr=ALL&start=&stop=&map=360&rs_term=&vtJerm=&speciesType=2&obj=qtl&fmt=5). *Fttflb*, and *Fpmoif10a* are two QTLs with similar phenotypes and map positions (labeled in black). *Ttda6, FcvfXa* and *FcvfXb* are three novel loci that do not overlap those listed on RGD (labeled in red). All the other 11 QTLs are overlap some known bone QTLs (usually BMD) but are now linked to specific microCT bone traits and are therefore new or refined (labeled in green).

**Table 1.**
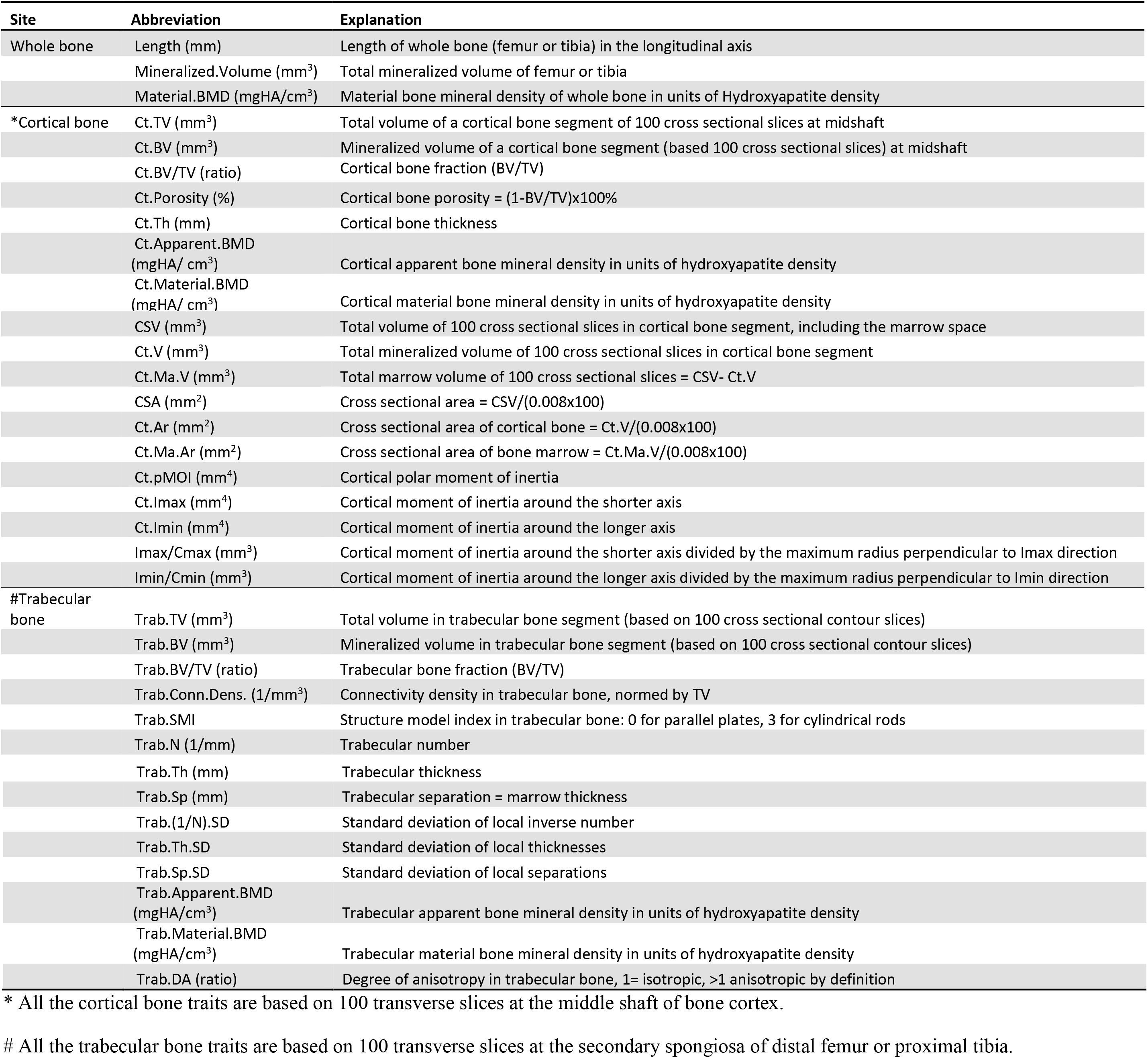
Abbreviations and explanations of bone traits measured by microCT

| Site | Abbreviation | Explanation |
| --- | --- | --- |
| Whole bone | Length (mm) | Length of whole bone (femur or tibia) in the longitudinal axis |
|  | Mineralized.Volume (mm<sup>3</sup>) | Total mineralized volume of femur or tibia |
|  | Material.BMD (mgHA/cm<sup>3</sup>) | Material bone mineral density of whole bone in units of Hydroxyapatite density |
| *Cortical bone | Ct.TV (mm<sup>3</sup>) | Total volume of a cortical bone segment of 100 cross sectional slices at midshaft |
|  | Ct.BV (mm<sup>3</sup>) | Mineralized volume of a cortical bone segment (based 100 cross sectional slices) at midshaft |
|  | Ct.BV/TV (ratio) | Cortical bone fraction (BV/TV) |
| | Ct.Porosity (%) | Cortical bone porosity = $(1 - BV/TV) \times 100\%$ |
|  | Ct.Th (mm) | Cortical bone thickness |
|  | Ct.Apparent.BMD (mgHA/ cm<sup>3</sup>) | Cortical apparent bone mineral density in units of hydroxyapatite density |
|  | Ct.Material.BMD (mgHA/ cm<sup>3</sup>) | Cortical material bone mineral density in units of hydroxyapatite density |
|  | CSV (mm<sup>3</sup>) | Total volume of 100 cross sectional slices in cortical bone segment, including the marrow space |
|  | Ct.V (mm<sup>3</sup>) | Total mineralized volume of 100 cross sectional slices in cortical bone segment |
|  | Ct.Ma.V (mm<sup>3</sup>) | Total marrow volume of 100 cross sectional slices = CSV- Ct.V |
| | CSA (mm<sup>2</sup>) | Cross sectional area = $CSV / (0.008 \times 100)$ |
| | Ct.Ar (mm<sup>2</sup>) | Cross sectional area of cortical bone = $Ct.V / (0.008 \times 100)$ |
| | Ct.Ma.Ar (mm<sup>2</sup>) | Cross sectional area of bone marrow = $Ct.Ma.V / (0.008 \times 100)$ |
|  | Ct.pMOI (mm<sup>4</sup>) | Cortical polar moment of inertia |
|  | Ct.Imax (mm<sup>4</sup>) | Cortical moment of inertia around the shorter axis |
|  | Ct.Imin (mm<sup>4</sup>) | Cortical moment of inertia around the longer axis |
|  | Imax/Cmax (mm<sup>3</sup>) | Cortical moment of inertia around the shorter axis divided by the maximum radius perpendicular to Imax direction |
|  | Imin/Cmin (mm<sup>3</sup>) | Cortical moment of inertia around the longer axis divided by the maximum radius perpendicular to Imin direction |
| #Trabecular bone | Trab.TV (mm<sup>3</sup>) | Total volume in trabecular bone segment (based on 100 cross sectional contour slices) |
|  | Trab.BV (mm<sup>3</sup>) | Mineralized volume in trabecular bone segment (based on 100 cross sectional contour slices) |
|  | Trab.BV/TV (ratio) | Trabecular bone fraction (BV/TV) |
|  | Trab.Conn.Dens. (1/mm<sup>3</sup>) | Connectivity density in trabecular bone, normed by TV |
|  | Trab.SMI | Structure model index in trabecular bone: 0 for parallel plates, 3 for cylindrical rods |
|  | Trab.N (1/mm) | Trabecular number |

|  |  |
| --- | --- |
| Trab.Th (mm) | Trabecular thickness |
| Trab.Sp (mm) | Trabecular separation = marrow thickness |
| Trab.(1/N).SD | Standard deviation of local inverse number |
| Trab.Th.SD | Standard deviation of local thicknesses |
| Trab.Sp.SD | Standard deviation of local separations |
| Trab.Apparent.BMD<br>(mgHA/cm<sup>3</sup>) | Trabecular apparent bone mineral density in units of hydroxyapatite density |
| Trab.Material.BMD<br>(mgHA/cm<sup>3</sup>) | Trabecular material bone mineral density in units of hydroxyapatite density |
| Trab.DA (ratio) | Degree of anisotropy in trabecular bone, 1= isotropic, >1 anisotropic by definition |

\* All the cortical bone traits are based on 100 transverse slices at the middle shaft of bone cortex.

### Statistical analysis

Bones were harvested immediately after sacrifice from roughly equal numbers of males and females ranging from 50 days to 375 days of age (100 ± 56 SD). This age range is equivalent to early adulthood to middle age in humans ^(25)^. Body weight ranged from 13.4 to 48.5 g (24.9 ± 5.1 SD). The relation between body weight and the logarithm of age (log age) fits a linear regression reasonably well in which weight (g) = −3.5 + 14.5(log10 of age), with an *r* = 0.45 and *p* < 0.0001.

All data and metadata on cases used in this study are provided in the **Supplemental Data S1** (*BXD_bone_data_master_table*). To minimize effects of age as a confounder, we performed linear regression for each variable across 307 male samples and 290 female samples separately, using logarithm of age as a predictor. Log age-corrected values in all tables for each trait and for each sex were computed by adding the residuals to means for male or female samples (**Supplemental Data S1**. Sheet: *Female_Raw_Res_Corr* and *Male_Raw_Res_Corr*). Since the average age of males and females was ~100 days, the corrected values by case, by strain, and by sex should be considered as those that will typically be measured at this age. Both the original values and the corrected values are provided in **Supplemental Data S1** (*BXD_bone_data_master_table*). Sex-averaged values were computed as above by fitting the cofactors sex and logarithm of age (without grouping by strain) across the entire data set. Means were added to the residuals, and these values were summarized to generate sex- and age-corrected strain means.

We also analyzed the relation between body weight and bone length/volume before and after correction for age. As expected, there is a strong positive correlation before age correction (body weight and bone volume covary with an *r* of 0.61, both sexes combined). After the log age correction, there is still a significant association between body weight and bone parameters. For example, the correlation between femur volume and body weight is 0.35. We chose not to correct for this source of variation since body weight and body size are also key variables of interest. But this does mean that bone data sets need to be considered in light of general variation in body size. All bone trait data used in subsequent analyses– heritability, trait covariance, and trait genetic mapping–include the adjustment for log age only. Effects of sex and strain as predictors were estimated by ANOVA.

After generating the corrected values we again searched for outliers at both the level of individual cases and strain means. All cases appear to be within normal limits when traits are examined individually. However there are strains such as BXD13 with small sample size (*n* = 2 males) and that are clearly outliers for some traits. These cases and data were censored in some analyses as described in *Results* and figure legends, either by complete removal or by winsorizing the outliers ^(26)^. Mean strain data entered into GeneNetwork has been reviewed and when necessary, has also been winsorized. However we do provide the original value in trait descriptions. Users can revert to the original data as needed.

The broad-sense heritability (*h*^2^) was calculated to estimate the effect of genetic factors on variance of bone traits. It is the genetic variance (variance among strain means) divided by the total variance (variance among all measurements). The variance and bias of the estimate of *h*^2^ was computed using a “drop-one-out” resampling jackknife procedure using JMP Pro 12 ^(27, 28)^. This involved calculating heritabilities for subsamples of data, each deleting all data for one strain (*h*_(-*i*)_). The jackknife variance is

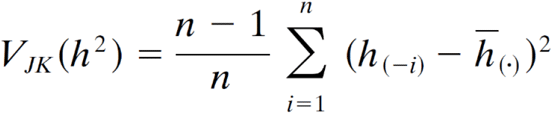

Where 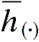 is the mean heritability of all strains.

To test if there is sex difference between female and male heritabilities, we computed Z value:

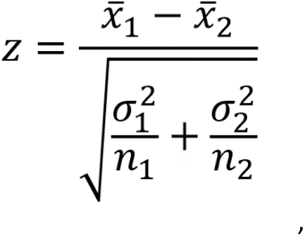

where 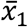, 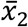 are the average of female and male heritabilities of each bone trait, *σ_1_^2^* and *σ_2_^2^* are jackknife variances of female and male heritabilities, respectively. *n_1_* = *n_2_* = 35. |*z*| ≥ 3.18 is considered significant with a Bonferroni corrected *p* < 0.00143 (0.05/35 ≌ 0.00143, two-tailed).

### Correlation analysis

We studied correlations among bone phenotypes using strain averages. We selected three representative traits for each of three major categories: whole bone (GeneNetwork Record ID 18130, 18131, 18132), cortical bone (GN 18134, 18136, 18141), and trabecular bone (GN 18146, 18148, 18149). Since the sample size is generally reasonably large (n ~60), we used Pearson product-moment correlations, and confirmed that results were not sensitive to outliers (BXD13 and BXD78 were often outliers for both sexes). We also correlated new bone phenotypes with 5000 previously published phenotypes for BXD family (17). Finally, we computed correlations between bone microtraits and an adult femur mRNA expression data (UCLA GSE27483 BXD Bone Femur ILM Mouse WG-6 v1.1 (Jan13) RSN that is included in GeneNetwork (GN accession number: GN410) generated by Farber and colleagues ^(14, 29)^. This data set includes expression values for 32 BXD strains and other strains for which we have matched bone phenotypes. Since the overlap of sample size is modest, we used rank order correlations in comparing bone phenotypes with expresssion data.

### QTL mapping

We carried out conventional interval mapping using Haley-Knott regression equations ^(30)^ as implemented in GeneNetwork. To estimate genome-wide thresholds of significance we permuted phenotypes 2000 or more times. Confidence intervals were defined as the chromosomal region within a 1.5 LOD (log of the odds) drop from the linkage peak. We have taken several approaches to evaluate the consistency of QTL results, including the use of two genotype files, different methods to correct for variation attributable to age and sex, and several different mapping algorithms that handle kinship relations. In total we mapped 50 traits for males, for females, and the sexes combined. Initial analysis was corrected for logarithm of age, but for comparison, we also remapped using only young animals within a relatively narrow age range. All aspects of this analysis can be reviewed and replicated using Record IDs in GeneNetwork (Table 4 and **Supplemental Data S1**).

#### Classic and new genotype files

We used two genotype files for mapping. The first file is an older genotype file that has been used by almost all investigators from 2002 through to late 2016. We refer to this as the “classic” genotype file because it has been used in hundreds of studies. The second file was released in January 2017 and includes roughly twice as many markers. Both files are available at www.genenetwork.org/webqtl/main.py?FormID=sharinginfo&GN_AccessionId=600. There is of course, a great deal of similarity between these files. The main difference is that several BXD strains were not fully inbred during the earlier phase of genotyping. Since the mice that we have studied here were born between 2011 and 2013, it is useful to compare results using both files.

#### Age effects

We were concerned that animals older than 150 days may have sufficiently different bone architecture that the age adjustment would not compensate fully. For this reason, we compared results based on the complete data set of 597 cases corrected for logAge (GN 18130 - 18279) to a subset of 484 cases ranging from 65 to 116 days and processed without any age correction (GN 18986 - 19086). This age range is equivalent to young adults in humans ^(25)^.

#### Mapping algorithms

Differences in algorithms and their sensitivities to trait distribution and kinship among strains will have effects on mapping results, in particular, on the maximum LRS scores. We therefore remapped traits–particular the 13 traits that gave strong initial results–using complementary algorithms that account for kinship and cofactors such as age and sex. These methods include several variants of R/qtl ^(31–33)^, and two algorithms that explicitly model kinship–pyLMM ^(34)^ and GEMMA ^(35, 36)^. All algorithms were run using implementations that are part of GeneNetwork 2 (gn2.genenetwork.org).

#### Composite interval mapping

Seven of 16 traits have QTLs on several chromosomes. For example, traits associated with femur moment of inertia (GN 18191, 18192, 18193) have loci on Chrs 7, 10, and 17 (*Fpmoif7, Fpmoif10a, Fpmoif10b* and *Fmoif17*). In cases with multiple QTLs, we wanted to ensure that the loci were in statistical linkage ^(37)^. To ensure independence, we selected background markers close to each QTL and remapped using composite interval mapping methods.

#### Interaction effects

While we have studied 60 or more BXD strains of both sexes, the sample size is marginal to test for two-way epistatic interactions. We used the pair-scan module in GeneNetwork and estimated empirical *p* values of the full model (V = Q1 + Q2 + Q1×Q2), where V is the trait variance, Q1 and Q2 are additive effect variance components for two QTLs, and Q1×Q2 is the variance attributable to an epistatic interaction. For these cases we reviewed results to ensure adequate distribution of genotypes among the four two-locus combinations of genotypes (*B/B, B/D, D/B*, and *D/D*), and to avoid effects generated by outliers. In general, we biased our pair-scan analysis is favor of results that included one or more loci with significant additive effects (LRS >10). We also tested for sex-by-QTL (S×Q) interactions using the model: V = Q1 + S + S×Q1. Sample size within strain was generally not sufficient to test for sex-by-strain differences.

### Gene ontology candidate analysis

To evaluate candidate genes using a more objective and unbiased method we exploited a femur gene expression data set (GN410) for the BXD strains ^(14)^. We used the following workflow (Figure 2):

1. We selected a list of 34 GO terms of eight major categories linked to bone and connective tissues (see **Supplemental Data S3**. Sheet: *GO genes*).
2. We extracted the list of genes linked to these 34 GO categories for mouse. Gene symbols were curated and edited to improve symbol alignment with the BXD femur gene expression database (14).
3. All the 46,621 Illumina probe sets corresponding to each key bone GO categories were used to validate the GO alignment using the femur mRNA expression data (GN410). For example for GO:0001503 (ossification) we expect each of the top 20 GO genes to generate lists of expression covariates that are themselves highly enriched in this category. As expected, the top 1,000 covariates of *Alpl* have an enrichment *p* of 10^-20^ for ossification. In the majority of cases this process using the bone expression data validate the precise parental GO classification.
4. Having established the validity of the adult femur mRNA expression data to detect GO enrichment for known bone-associated genes in step 3, we applied the same procedures to every candidate gene by computing its enrichment for bone GO terms. To be considered a strong biological candidate, these genes had to have a GO enrichment equal to or greater than the average of known members of the ontology. For example, genes known to be associated with GO: 0001503 have an average GO enrichment *p* value of 10^-8^—a value that candidates had to match or beat exceed.

**Figure 2.**
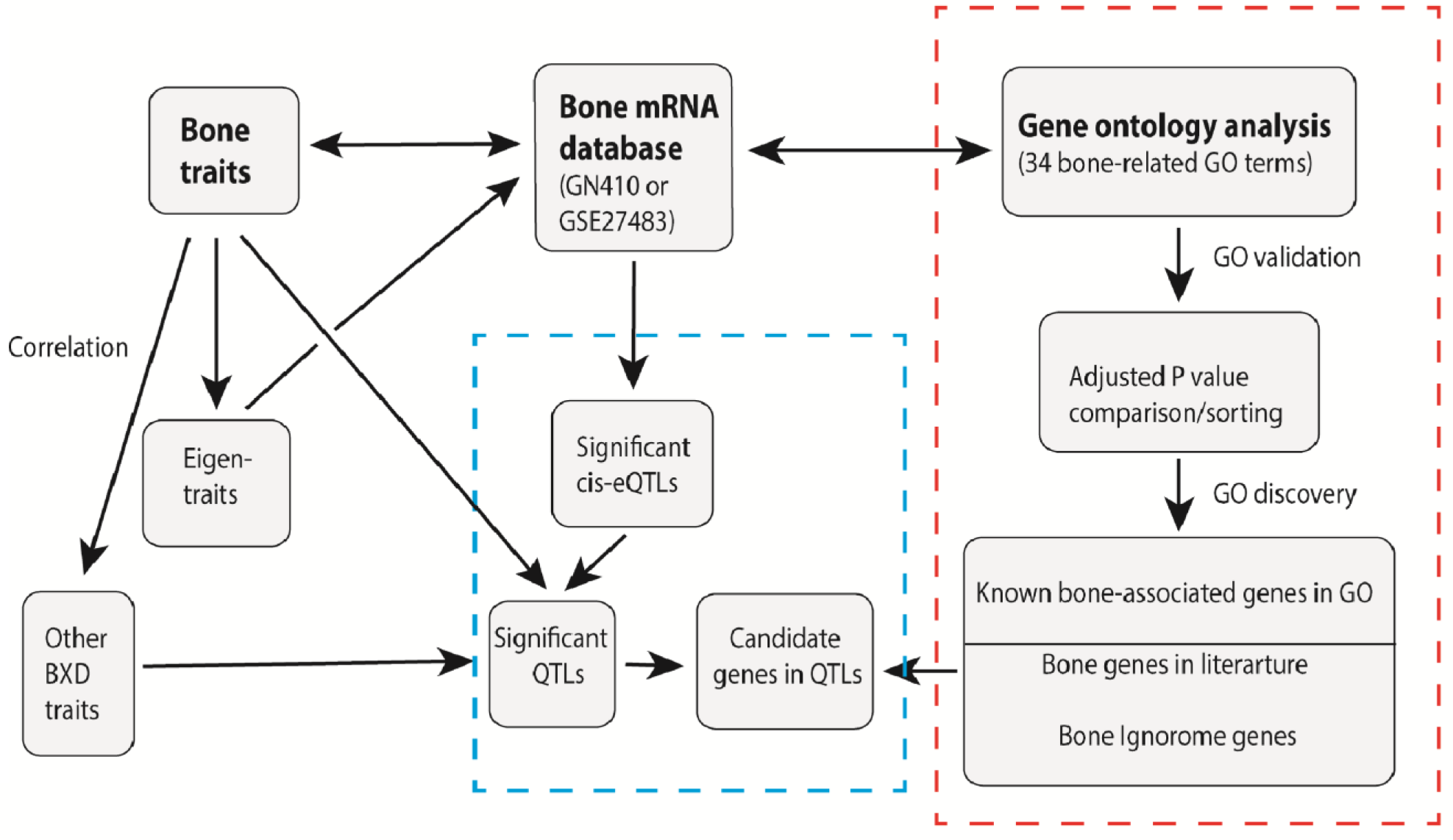
QTL-candidate gene analysis workflow. We exploited three primary data sources (bold font at top of figure) for this analysis. **Bone traits** were correlated to all **Other BXD traits** in GN. We also conducted PCA with the attempt to summarize all 150 bone traits into **Eigentraits** (six eigentraits were listed from GN 18424 to GN 18429). Finally, **Bone traits** were compared against a comprehensive **Bone mRNA database** GN410. The QTL analysis section (box with blue dashed line, bottom center) consists of three boxes that lead to **Candidate genes in QTLs**. The GO analysis section (box with red dashed line to right) summarizes the method used to generate “bone scores” for all probes in the **Bone mRNA database**.

### Scoring system of candidate genes

As shown in the *Results*, we have mapped 16 QTLs that typically contain 45 to 174 positional candidate genes within the 1.5 LOD confidence intervals. Collectively, these 16 QTLs include 1,638 protein-coding genes. Based on the assignment of these candidates to 34 bone GO terms (e.g.,” bone development”, “ossification”, “bone remodeling”, etc.), only 2% (*n* = 36 genes) have been associated with bone biology in the literature. We wanted to develop a more comprehensive and objective method to evaluate the remaining 98% of candidate genes for possible association with bone biology.

To do this we adapted methods used in previous studies to define potential candidate genes by computing so-called *bone scores* using methods similar to those described in references ^(22, 23, 38)^. In our case, we specifically defined a bone score that estimates the potential association between each gene and a reference set of 770 genes already associated with 34 bone GO terms. We exploited the same large femur expression dataset (GN410) that includes probes that target essentially all protein-coding transcripts. For each of 46,621probes, we first calculated the absolute Spearman correlation between the mRNA abundance of this probe and all other probes. We then selected the top 1,000 probes with the highest correlations and performed a GO enrichment analysis (biological process) based on the hypergeometric test ^(39–41)^:

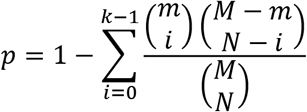

where *M* represents the total number of genes targeted by all probe IDs (*n* = 30,880); *N* represents the number of unique genes among the top 1,000 covariates; *m* represents the number of genes listed in the GO term; and *k* represents the number of genes among the top 1,000 covariates that are in the GO term.

For example, the top 1,000 covariates of *Alpl* (alkaline phosphatase, probe ID ILM2340168) includes 369 genes associated with “ossification” (G0:0001503). Even after correction for multiple comparisons, the *p* value of this GO term enrichment is 5×10^-23^. We computed enrichment *p* values for a set of 34 bone-associated GO terms (**Supplemental Data S3**). The average *p* value of 40 well known ossification genes such as *Alpl* was used as reference standard against which we compared all other genes/probes (also see **Supplemental Data S3**). Many positional candidate genes had *p* values that were as good as or better than those of these 36 known bone-associated genes. We converted values to −log_10_ (*p*) across all 34 terms and used the average value as a GO-associated “**bone score**”. Genes such as *Alpl* that are linked to several bone GO terms typically have average scores above 1 (the peak score is 10.5 for *Col15a1*). Genes linked to 10 or more terms typically have scores above 2. Genes without links to bone have averages well below 1. A large subset of genes were further defined as ***bone ignorome*** genes– those that have high bone scores based on the GO analysis but no known literature associated with the skeletal system.

Finally, we generated a **summary candidate score** on a scale of 1 to 10 points for all genes based on:

1. **Average bone score (1 to 3)**. The gene is grading-scored 1 if its average bone score across 34 GOs is between 0 and 1; 2 if between 1 and 3; and scored 3 if greater than 3.
2. **Highest bone score (0 to 1)**. Some genes are restricted to only one or two bone GO, with high bone score in these a few Go, but very low average score across 34 GOs. We therefore assigned these genes a grading score of 1 provided that its highest bone score is between 5 and 10; and a grading score of 2 if greater than 10. None of 1,638 positional genes has a highest bone score greater than 10.
3. **Coding DNA variants (0 to 3)**. In a recent study we defined 35,068 coding SNPs in the BXDs, of which 11,979 are non-synonymous (nsSNP) with Grantham scores (complete list is Supplementary Data 4, of Wang et al ^(17)^). Grantham scores estimate the impact of non-synonymous variants into classes based on chemical dissimilarity of amino acids ^(42)^. A higher Grantham score reflects a greater potential impact ^(43)^. The gene is grading-scored 1 if it has a sum of Grantham values less than 100; 2 if the sum is between 100 and 300; and 3 if the sum is greater than 300. We also identified 173 SNPs with nonsense, frame shift, or splice site mutation (Supplementary Data 5 of Wang et al. ^(17)^). Genes with these variants were scored 3.
4. **Cis-regulation (0 to 3)**. If a gene is cis-regulated in bone, it receives a score of 3. If in cartilage and muscle, it is scored 2. If in other tissue, it is scored 1.

All genes received a **summary candidate score** on a 1 to 10 point scale (**Supplemental Data S4** and Table 5). We focused most analysis on a subset of 212 candidates within 16 QTLs with scores greater than 4. With the goal of functional validation of candidates, we needed to shorten even this list even more, and we therefore restricted analysis to the top 50 candidates–the top 10–20 from seven robust QTLs.

### Candidate gene analysis using RGD, GWAS, and KO resources

We scored candidates using several public resources:

1. The Rat Genome Database (RGD, www.rgd.mcw.edu). This resource provides current genome and phenome data for mouse, rat, and human ^(44)^. We downloaded a list of 1,000 genes from RGD using “bone” as the keyword in all species and used this set to decide if candidate genes have known bone association.
2. Human GWAS gene compendium (www.ebi.ac.uk/gwas) using “bone” and “skeletal” as keywords at a *p* threshold ≤ 5 × 10^-4^. We download a list of 1125 genes associated with diseases that impact bone and skeletal system.
3. International Mouse Phenotyping Consortium, a collection of phenotype data on mouse knockout lines (IMPC, www.mousephenotype.org). We searched and downloaded a list of 699 genes associated with abnormal skeletal phenotype (mouse phenotype MP: 0005508).

### Candidate genes underlying human GWAS loci for BMD

Using the 2017 UKBB eBMD GWAS data ^(45)^, we defined bins between the furthest upstream and downstream SNPs with a linkage disequilibrium *r^2^* of ≥ 0.7 with the lead SNP, and calculated from European populations in the 1000 Genomes Phase III data ^(46)^. For each bin, we identified all genes that overlap a bin. If no genes intersected the bin, the nearest upstream and downstream genes were included. This yielded 731 genes underlying GWAS loci for eBMD (**Supplemental Data S4**, sheet “731_BMD_GWAS_genes”).

## Results

We measured 25 microCT phenotypes from femur and tibia in a total of 63 BXD strains. Gross bone morphology and size vary significantly by age and sex. The range is a function of dimensionality, and as expected volumetric measurements typically have a wider range compared to linear and areal measurements. For example, the maximum range of femoral length among individuals is ~1.55-fold—from 10.9 to 17.0 mm, while that for femoral volume is ~3-fold—from 8.6 to 26.2 mm3. Material BMD has comparatively low variation of 1.3-fold.

The coefficient of variance (CV, SD/mean x 100) is a better metric for comparing variation among traits. CVs will be a function of biological variation, developmental variation (e.g. right versus left differences), technical error, and even the dimensionality of the measure. For example the CV of material BMD range from 2.9% to 4.1% (3.7% ± 0.1%, both sexes combined, including material BMD of whole bone, cortical and trabecular bone compartments). This is also true for other ratio-based traits, including cortical bone fraction (0.3 – 0.4%) and trabecular degree of anisotropy (9.7 – 11.1%). In contrast, several clusters of related bone microtraits have high CVs. For example, several parameters related to cortical moment of inertia are sensitive to slight differences in geometry and have CVs that are 100-fold higher—from 33 to 51% in both sexes and both bones. At trabecular site the variation is particularly high because of complex microstructure and active bone turnover and comparatively high technical and sampling error. CVs in this cluster of traits also range from 33 to 54%, including trabecular volume, bone fraction, and connectivity density.

### BXD strain averages

Much of the variation among mice is attributable to differences among the inbred BXD strains, and in the following sections we discuss correlations and mapping results of strain means. All strain averages of nine representative femoral phenotypes, split by sex and corrected for age, are provided in Table 2, including three for whole bone (bone length, mineralized volume and material BMD); three for cortical bone (cortical thickness, porosity and polar moment inertia); three for trabecular bone (trabecular thickness, bone fraction and connectivity density). Averages for the other 16 femur and 25 tibia phenotypes can be computed from data in **Supplemental Data S1**, but are also presented in GeneNetwork (GN IDs 18130 to 18279) for sex averages, females alone, and males alone.

**Table 2.**
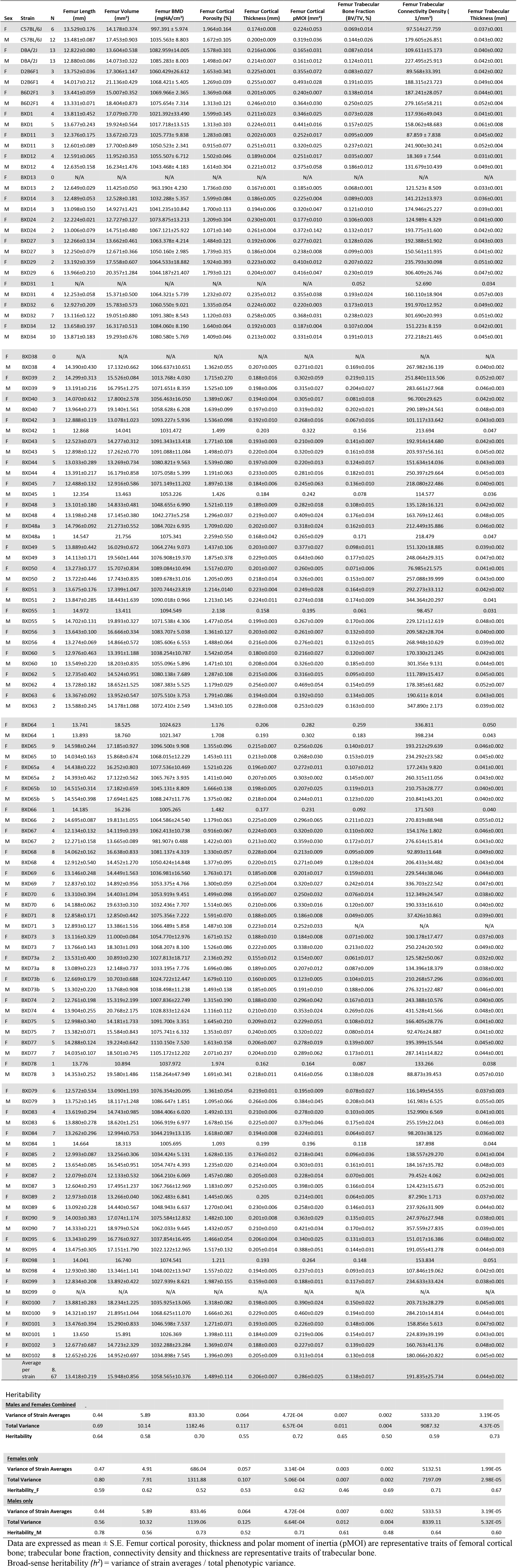
Summary of Femur Phenotypes with Heritability

| Sex | Strain | N | Femur Length (mm) | Femur Volume (mm<sup>3</sup>) | Femur BMD (mgHA/cm<sup>3</sup>) | Femur Cortical Porosity (%) | Femur Cortical Thickness (mm) | Femur Cortical pMOI (mm<sup>4</sup>) | Femur Trabecular Bone Fraction (BV/TV, %) | Femur Trabecular Connectivity Density (1/mm<sup>3</sup>) | Femur Trabecular Thickness (mm) |
| --- | --- | --- | --- | --- | --- | --- | --- | --- | --- | --- | --- |
| F | C57BL/6J | 6 | 13.529±0.176 | 14.178±0.374 | 997.391 ± 5.974 | 1.964±0.164 | 0.174±0.008 | 0.224±0.053 | 0.069±0.014 | 97.514±27.759 | 0.037±0.001 |
| M | C57BL/6J | 12 | 13.481±0.087 | 17.453±0.903 | 1035.563± 8.803 | 1.672±0.105 | 0.200±0.009 | 0.319±0.036 | 0.144±0.026 | 179.605±26.851 | 0.043±0.002 |
| F | DBA/2J | 13 | 12.822±0.080 | 13.604±0.538 | 1082.959±14.005 | 1.578±0.101 | 0.216±0.006 | 0.165±0.031 | 0.087±0.014 | 109.611±15.173 | 0.040±0.002 |
| M | DBA/2J | 13 | 12.880±0.086 | 14.073±0.322 | 1085.283± 8.003 | 1.498±0.047 | 0.214±0.007 | 0.161±0.012 | 0.124±0.011 | 227.495±25.913 | 0.042±0.001 |
| F | D2B6F1 | 3 | 13.752±0.036 | 17.306±1.147 | 1060.429±26.612 | 1.653±0.341 | 0.225±0.001 | 0.355±0.072 | 0.083±0.027 | 89.568±33.391 | 0.042±0.002 |
| M | D2B6F1 | 4 | 14.017±0.212 | 21.136±0.429 | 1068.421± 5.405 | 1.269±0.039 | 0.255±0.007 | 0.493±0.028 | 0.191±0.035 | 188.315±23.723 | 0.049±0.004 |
| F | B6D2F1 | 3 | 13.441±0.059 | 15.007±0.352 | 1069.966± 2.365 | 1.369±0.068 | 0.201±0.005 | 0.240±0.007 | 0.138±0.014 | 187.241±28.057 | 0.044±0.001 |
| M | B6D2F1 | 4 | 13.331±0.071 | 18.404±0.873 | 1075.654± 7.314 | 1.313±0.121 | 0.246±0.010 | 0.364±0.030 | 0.250±0.025 | 279.165±58.211 | 0.052±0.004 |
| F | BXD1 | 4 | 13.811±0.452 | 17.079±0.770 | 1021.392±33.490 | 1.599±0.145 | 0.211±0.023 | 0.346±0.025 | 0.073±0.028 | 117.936±49.043 | 0.041±0.001 |
| M | BXD1 | 5 | 13.677±0.243 | 19.924±0.564 | 1017.718±13.515 | 1.313±0.103 | 0.224±0.011 | 0.441±0.016 | 0.157±0.025 | 158.062±48.683 | 0.061±0.008 |
| F | BXD11 | 3 | 12.376±0.175 | 13.672±0.723 | 1025.773± 9.838 | 1.283±0.081 | 0.202±0.003 | 0.252±0.017 | 0.095±0.009 | 87.859 ± 7.838 | 0.045±0.002 |
| M | BXD11 | 3 | 12.601±0.089 | 17.700±0.849 | 1050.523± 2.341 | 0.915±0.077 | 0.251±0.011 | 0.320±0.025 | 0.237±0.021 | 241.900±30.241 | 0.052±0.004 |
| F | BXD12 | 4 | 12.591±0.065 | 11.952±0.353 | 1055.507± 6.712 | 1.502±0.046 | 0.189±0.004 | 0.251±0.017 | 0.035±0.007 | 18.369 ± 7.544 | 0.031±0.001 |
| M | BXD12 | 4 | 12.635±0.158 | 16.234±1.476 | 1043.468± 4.183 | 1.614±0.304 | 0.221±0.012 | 0.375±0.058 | 0.186±0.012 | 131.679±10.439 | 0.049±0.001 |
| F | BXD13 | 0 | N/A | N/A | N/A | N/A | N/A | N/A | N/A | N/A | N/A |
| M | BXD13 | 2 | 12.649±0.029 | 11.425±0.050 | 963.190± 4.230 | 1.736±0.030 | 0.167±0.001 | 0.185±0.005 | 0.068±0.001 | 121.523± 8.509 | 0.033±0.001 |
| F | BXD14 | 3 | 12.489±0.053 | 12.528±0.181 | 1032.288± 5.357 | 1.599±0.084 | 0.186±0.005 | 0.225±0.004 | 0.089±0.003 | 141.212±13.973 | 0.036±0.001 |
| M | BXD14 | 3 | 13.098±0.150 | 14.927±1.421 | 1041.235±10.842 | 1.700±0.113 | 0.194±0.006 | 0.320±0.047 | 0.121±0.010 | 174.946±25.227 | 0.039±0.001 |
| F | BXD24 | 2 | 12.224±0.021 | 12.727±0.127 | 1073.875±13.213 | 1.209±0.104 | 0.230±0.001 | 0.177±0.010 | 0.106±0.003 | 124.989± 4.329 | 0.041±0.000 |
| M | BXD24 | 2 | 13.006±0.079 | 14.751±0.480 | 1067.121±25.922 | 1.071±0.140 | 0.261±0.004 | 0.372±0.142 | 0.132±0.017 | 193.775±31.600 | 0.042±0.002 |
| F | BXD27 | 3 | 12.266±0.134 | 13.662±0.461 | 1063.378± 4.214 | 1.484±0.121 | 0.192±0.006 | 0.277±0.021 | 0.128±0.026 | 192.388±51.902 | 0.043±0.003 |
| M | BXD27 | 3 | 12.250±0.079 | 12.671±0.366 | 1050.160± 2.985 | 1.739±0.315 | 0.186±0.004 | 0.238±0.008 | 0.099±0.003 | 150.561±11.935 | 0.041±0.002 |
| F | BXD29 | 2 | 13.192±0.359 | 17.558±0.607 | 1064.533±18.882 | 1.924±0.393 | 0.223±0.002 | 0.410±0.012 | 0.207±0.022 | 235.793±30.098 | 0.051±0.002 |
| M | BXD29 | 6 | 13.966±0.210 | 20.357±1.284 | 1044.187±21.407 | 1.793±0.121 | 0.204±0.007 | 0.416±0.047 | 0.230±0.019 | 306.409±26.746 | 0.047±0.002 |
| F | BXD31 | 1 | N/A | N/A | N/A | N/A | N/A | N/A | 0.052 | 52.690 | 0.034 |
| M | BXD31 | 4 | 12.253±0.058 | 15.371±0.500 | 1064.321± 5.739 | 1.232±0.072 | 0.235±0.012 | 0.355±0.038 | 0.193±0.024 | 160.110±18.904 | 0.057±0.003 |
| F | BXD32 | 6 | 12.927±0.209 | 15.783±0.573 | 1060.550± 9.021 | 1.335±0.054 | 0.224±0.002 | 0.220±0.003 | 0.173±0.013 | 191.970±12.952 | 0.049±0.002 |
| M | BXD32 | 7 | 13.116±0.122 | 19.051±0.880 | 1091.380± 8.543 | 1.120±0.033 | 0.258±0.005 | 0.368±0.031 | 0.238±0.023 | 301.690±20.993 | 0.051±0.002 |
| F | BXD34 | 12 | 13.658±0.197 | 16.317±0.513 | 1084.060± 8.190 | 1.640±0.064 | 0.192±0.003 | 0.187±0.004 | 0.107±0.004 | 151.223± 8.159 | 0.042±0.001 |
| M | BXD34 | 10 | 13.871±0.183 | 19.293±0.676 | 1080.580± 5.769 | 1.409±0.046 | 0.213±0.002 | 0.331±0.014 | 0.191±0.013 | 272.218±21.465 | 0.045±0.001 |
| F | BXD38 | 0 | N/A | N/A | N/A | N/A | N/A | N/A | N/A | N/A | N/A |
| M | BXD38 | 4 | 14.390±0.430 | 17.132±0.662 | 1066.637±10.651 | 1.362±0.055 | 0.207±0.005 | 0.271±0.021 | 0.169±0.016 | 267.982±36.139 | 0.040±0.002 |
| F | BXD39 | 2 | 14.299±0.313 | 15.526±0.084 | 1013.768± 4.030 | 1.715±0.270 | 0.188±0.016 | 0.302±0.059 | 0.219±0.115 | 251.840±113.506 | 0.052±0.007 |
| M | BXD39 | 9 | 13.191±0.216 | 16.795±1.275 | 1071.651± 8.359 | 1.525±0.109 | 0.198±0.006 | 0.315±0.027 | 0.204±0.027 | 283.661±27.968 | 0.046±0.003 |
| F | BXD40 | 3 | 14.070±0.612 | 17.800±2.578 | 1056.463±16.050 | 1.389±0.067 | 0.194±0.004 | 0.305±0.017 | 0.081±0.018 | 96.700±29.625 | 0.042±0.002 |
| M | BXD40 | 7 | 13.964±0.273 | 19.140±1.561 | 1058.628± 6.208 | 1.639±0.099 | 0.197±0.010 | 0.319±0.032 | 0.202±0.021 | 290.189±24.561 | 0.048±0.003 |
| F | BXD42 | 3 | 12.888±0.119 | 13.078±1.023 | 1093.227± 5.936 | 1.536±0.098 | 0.192±0.010 | 0.268±0.016 | 0.067±0.016 | 101.117±33.642 | 0.043±0.003 |
| M | BXD42 | 1 | 12.868 | 14.041 | 1031.472 | 1.499 | 0.203 | 0.322 | 0.156 | 213.694 | 0.047 |
| F | BXD43 | 5 | 12.523±0.073 | 14.277±0.312 | 1091.343±13.418 | 1.771±0.108 | 0.193±0.003 | 0.210±0.009 | 0.141±0.007 | 192.914±14.680 | 0.042±0.001 |
| M | BXD43 | 5 | 12.898±0.122 | 17.262±0.770 | 1091.088±11.084 | 1.498±0.073 | 0.220±0.004 | 0.320±0.029 | 0.161±0.038 | 203.937±56.161 | 0.045±0.002 |
| F | BXD44 | 5 | 13.033±0.289 | 13.269±0.734 | 1080.821± 9.563 | 1.539±0.080 | 0.197±0.009 | 0.220±0.013 | 0.124±0.017 | 151.634±14.036 | 0.043±0.003 |
| M | BXD44 | 4 | 13.391±0.217 | 16.179±0.858 | 1075.058± 5.399 | 1.191±0.063 | 0.233±0.005 | 0.281±0.016 | 0.182±0.031 | 250.397±29.664 | 0.045±0.003 |
| F | BXD45 | 7 | 12.488±0.132 | 12.916±0.586 | 1071.149±11.202 | 1.897±0.138 | 0.184±0.006 | 0.245±0.063 | 0.136±0.010 | 218.080±22.486 | 0.040±0.001 |
| M | BXD45 | 1 | 12.354 | 13.463 | 1053.226 | 1.426 | 0.184 | 0.242 | 0.078 | 114.577 | 0.036 |
| F | BXD48 | 3 | 13.101±0.180 | 14.833±0.481 | 1048.655± 6.990 | 1.521±0.119 | 0.189±0.009 | 0.282±0.018 | 0.108±0.015 | 135.128±16.121 | 0.042±0.002 |
| M | BXD48 | 4 | 13.198±0.248 | 17.145±0.380 | 1042.273±5.258 | 1.296±0.037 | 0.219±0.007 | 0.409±0.024 | 0.176±0.034 | 163.769±12.461 | 0.048±0.005 |
| F | BXD48a | 3 | 14.796±0.092 | 21.273±0.552 | 1084.702± 6.935 | 1.709±0.020 | 0.202±0.007 | 0.318±0.024 | 0.162±0.013 | 212.449±35.886 | 0.046±0.002 |
| M | BXD48a | 1 | 14.547 | 21.756 | 1075.341 | 2.259±0.550 | 0.168±0.042 | 0.265±0.029 | 0.171 | 218.479 | 0.047 |
| F | BXD49 | 5 | 13.889±0.442 | 16.029±0.672 | 1064.274± 9.073 | 1.437±0.106 | 0.203±0.007 | 0.377±0.027 | 0.098±0.011 | 151.320±18.885 | 0.039±0.002 |
| M | BXD49 | 3 | 14.113±0.171 | 19.560±1.444 | 1076.908±19.370 | 1.875±0.378 | 0.229±0.005 | 0.643±0.060 | 0.177±0.025 | 248.064±29.315 | 0.047±0.002 |
| F | BXD50 | 4 | 13.273±0.177 | 15.707±0.834 | 1089.084±10.494 | 1.517±0.070 | 0.201±0.007 | 0.260±0.005 | 0.071±0.006 | 76.985±21.575 | 0.041±0.001 |
| M | BXD50 | 2 | 13.722±0.446 | 17.743±0.835 | 1089.678±31.016 | 1.205±0.093 | 0.218±0.014 | 0.326±0.001 | 0.153±0.007 | 257.088±39.999 | 0.043±0.000 |
| F | BXD51 | 3 | 13.675±0.176 | 17.399±1.047 | 1070.744±23.819 | 1.214±.0140 | 0.223±0.004 | 0.249±0.028 | 0.164±0.019 | 292.273±33.112 | 0.042±0.002 |
| M | BXD51 | 2 | 13.847±0.285 | 18.443±1.639 | 1090.018± 0.966 | 1.213±0.145 | 0.224±0.011 | 0.274±0.038 | 0.174±0.009 | 344.364±20.297 | 0.041 |
| F | BXD55 | 1 | 14.972 | 13.411 | 1094.549 | 2.138 | 0.158 | 0.195 | 0.061 | 98.457 | 0.031 |
| M | BXD55 | 5 | 14.702±0.131 | 19.893±0.327 | 1071.538± 4.306 | 1.477±0.054 | 0.199±0.003 | 0.267±0.009 | 0.170±0.006 | 229.121±12.619 | 0.048±0.001 |
| F | BXD56 | 3 | 13.643±0.100 | 16.666±0.334 | 1083.707± 5.038 | 1.361±0.127 | 0.203±0.002 | 0.261±0.007 | 0.132±0.010 | 209.582±28.704 | 0.040±0.000 |
| M | BXD56 | 4 | 13.274±0.069 | 14.866±0.572 | 1085.606± 6.553 | 1.488±0.064 | 0.216±0.006 | 0.276±0.021 | 0.132±0.015 | 268.948±10.629 | 0.039±0.002 |
| F | BXD60 | 5 | 12.976±0.463 | 13.391±1.188 | 1038.254±10.787 | 1.542±0.054 | 0.180±0.010 | 0.216±0.027 | 0.120±0.007 | 170.330±21.245 | 0.042±0.001 |
| M | BXD60 | 10 | 13.549±0.220 | 18.203±0.835 | 1055.096± 5.896 | 1.471±0.101 | 0.208±0.004 | 0.326±0.020 | 0.185±0.010 | 301.356± 9.131 | 0.044±0.001 |
| F | BXD62 | 5 | 12.735±0.402 | 14.524±0.951 | 1080.138± 7.689 | 1.287±0.108 | 0.215±0.006 | 0.316±0.015 | 0.095±0.010 | 111.789±15.417 | 0.045±0.001 |
| M | BXD62 | 4 | 13.728±0.182 | 18.652±1.525 | 1087.383± 5.525 | 1.179±0.029 | 0.256±0.007 | 0.469±0.054 | 0.154±0.059 | 178.385±61.682 | 0.052±0.007 |
| F | BXD63 | 6 | 13.367±0.092 | 13.952±0.547 | 1075.510± 3.753 | 1.791±0.086 | 0.194±0.004 | 0.192±0.010 | 0.134±0.005 | 190.611± 8.014 | 0.043±0.001 |
| M | BXD63 | 2 | 13.588±0.245 | 14.178±1.088 | 1072.410± 2.549 | 1.343±0.105 | 0.228±0.008 | 0.253±0.029 | 0.163±0.010 | 347.890± 2.173 | 0.039±0.002 |
| F | BXD64 | 1 | 13.741 | 18.525 | 1024.623 | 1.176 | 0.206 | 0.282 | 0.259 | 336.811 | 0.050 |
| M | BXD64 | 1 | 13.893 | 18.760 | 1021.347 | 1.708 | 0.193 | 0.302 | 0.183 | 398.234 | 0.043 |
| F | BXD65 | 9 | 14.598±0.244 | 17.185±0.927 | 1096.500± 9.908 | 1.355±0.096 | 0.215±0.007 | 0.256±0.026 | 0.140±0.017 | 193.212±29.639 | 0.046±0.002 |
| M | BXD65 | 10 | 14.034±0.163 | 15.868±0.674 | 1068.015±12.229 | 1.453±0.111 | 0.213±0.008 | 0.268±0.030 | 0.153±0.019 | 234.292±23.582 | 0.045±0.002 |
| F | BXD65a | 4 | 14.438±0.222 | 16.252±0.803 | 1077.536±10.469 | 1.521±0.226 | 0.196±0.007 | 0.272±0.011 | 0.107±0.012 | 177.243± 9.820 | 0.041±0.001 |
| M | BXD65a | 2 | 14.393±0.462 | 17.122±0.562 | 1065.767± 3.935 | 1.411±0.040 | 0.207±0.005 | 0.303±0.002 | 0.145±0.007 | 260.315±11.056 | 0.043±0.002 |
| F | BXD65b | 10 | 14.515±0.314 | 17.182±0.659 | 1045.131± 8.809 | 1.666±0.138 | 0.198±0.005 | 0.207±0.025 | 0.119±0.013 | 210.753±28.777 | 0.040±0.001 |
| M | BXD65b | 5 | 14.554±0.398 | 17.694±1.625 | 1088.247±11.776 | 1.375±0.082 | 0.218±0.004 | 0.244±0.011 | 0.123±0.020 | 210.841±43.201 | 0.040±0.002 |
| F | BXD66 | 1 | 14.185 | 16.236 | 1005.265 | 1.482 | 0.177 | 0.231 | 0.092 | 171.503 | 0.040 |
| M | BXD66 | 2 | 14.695±0.087 | 19.813±1.055 | 1064.586±24.540 | 1.179±0.063 | 0.225±0.009 | 0.296±0.065 | 0.211±0.023 | 270.819±88.948 | 0.055±0.012 |
| F | BXD67 | 4 | 12.134±0.132 | 14.119±0.193 | 1062.413±10.738 | 0.916±0.067 | 0.224±0.003 | 0.320±0.020 | 0.110±0.002 | 154.176± 1.802 | 0.046±0.001 |
| M | BXD67 | 2 | 12.271±0.158 | 13.665±0.089 | 981.907± 0.488 | 1.422±0.003 | 0.213±0.002 | 0.359±0.030 | 0.172±0.017 | 276.614±15.814 | 0.043±0.002 |
| F | BXD68 | 8 | 14.062±0.162 | 16.638±0.833 | 1081.137± 4.319 | 1.330±0.057 | 0.228±0.004 | 0.213±0.009 | 0.095±0.009 | 92.893±11.648 | 0.049±0.002 |
| M | BXD68 | 4 | 12.912±0.540 | 14.452±1.270 | 1050.424±14.848 | 1.377±0.095 | 0.220±0.015 | 0.271±0.049 | 0.128±0.024 | 206.433±34.482 | 0.043±0.004 |
| F | BXD69 | 6 | 13.146±0.248 | 14.449±1.563 | 1036.981±16.560 | 1.763±0.171 | 0.185±0.008 | 0.201±0.017 | 0.159±0.031 | 229.544±38.046 | 0.044±0.003 |
| M | BXD69 | 7 | 12.837±0.102 | 14.892±0.956 | 1053.375± 4.766 | 1.300±0.059 | 0.225±0.004 | 0.320±0.027 | 0.242±0.014 | 336.703±22.542 | 0.047±0.001 |
| F | BXD70 | 6 | 13.310±0.394 | 14.403±1.094 | 1053.919± 9.451 | 1.499±0.098 | 0.195±0.007 | 0.250±0.032 | 0.076±0.014 | 112.349±24.547 | 0.038±0.002 |
| M | BXD70 | 6 | 14.188±0.062 | 19.633±0.310 | 1032.436± 7.707 | 1.514±0.065 | 0.210±0.006 | 0.330±0.016 | 0.120±0.007 | 190.333±16.610 | 0.040±0.002 |
| F | BXD71 | 8 | 12.858±0.171 | 12.850±0.442 | 1075.356± 7.222 | 1.591±0.070 | 0.188±0.005 | 0.186±0.008 | 0.049±0.005 | 37.426±10.861 | 0.039±0.001 |
| M | BXD71 | 3 | 12.893±0.127 | 13.386±1.516 | 1066.489± 5.858 | 1.487±0.108 | 0.223±0.014 | 0.252±0.033 | N/A | N/A | N/A |
| F | BXD73 | 3 | 13.116±0.329 | 11.000±0.084 | 1054.770±12.976 | 1.671±0.152 | 0.188±0.010 | 0.184±0.008 | 0.071±0.002 | 100.178±17.477 | 0.037±0.003 |
| M | BXD73 | 7 | 13.766±0.143 | 18.303±1.093 | 1068.207± 8.100 | 1.526±0.086 | 0.222±0.005 | 0.338±0.020 | 0.213±0.022 | 250.224±20.592 | 0.049±0.002 |
| F | BXD73a | 2 | 13.531±0.400 | 10.893±0.230 | 1027.813±18.717 | 2.136±0.292 | 0.155±0.012 | 0.154±0.007 | 0.061±0.017 | 125.582±50.067 | 0.032±0.002 |
| M | BXD73a | 8 | 13.089±0.223 | 12.148±0.737 | 1033.195± 7.776 | 1.696±0.086 | 0.189±0.005 | 0.207±0.012 | 0.087±0.009 | 134.396±18.379 | 0.038±0.002 |
| F | BXD73b | 6 | 12.669±0.179 | 10.703±0.688 | 1024.722±12.447 | 1.679±0.110 | 0.160±0.005 | 0.123±0.005 | 0.104±0.015 | 210.268±57.296 | 0.036±0.001 |
| M | BXD73b | 5 | 13.302±0.220 | 13.768±0.908 | 1038.498±11.238 | 1.493±0.138 | 0.185±0.005 | 0.191±0.010 | 0.188±0.006 | 276.321±22.487 | 0.046±0.001 |
| F | BXD74 | 2 | 12.761±0.198 | 15.319±2.199 | 1007.836±22.749 | 1.315±0.190 | 0.188±0.030 | 0.296±0.042 | 0.167±0.013 | 243.388±10.576 | 0.040±0.005 |
| M | BXD74 | 4 | 13.904±0.255 | 20.768±2.175 | 1028.833±12.624 | 1.116±0.112 | 0.210±0.010 | 0.353±0.024 | 0.269±0.026 | 431.528±41.566 | 0.048±0.001 |
| F | BXD75 | 5 | 12.998±0.340 | 14.181±1.733 | 1091.700± 3.351 | 1.645±0.210 | 0.209±0.012 | 0.229±0.051 | 0.108±0.012 | 166.405±28.776 | 0.041±0.002 |
| M | BXD75 | 7 | 13.382±0.071 | 15.584±0.843 | 1075.741± 6.332 | 1.353±0.037 | 0.240±0.005 | 0.320±0.022 | 0.080±0.014 | 92.476±24.887 | 0.041±0.002 |
| F | BXD77 | 5 | 14.288±0.124 | 19.224±0.642 | 1110.150± 7.520 | 1.613±0.158 | 0.206±0.007 | 0.278±0.019 | 0.139±0.007 | 195.399±15.544 | 0.045±0.002 |
| M | BXD77 | 7 | 14.035±0.107 | 18.501±0.745 | 1105.172±12.202 | 2.071±0.237 | 0.204±0.010 | 0.289±0.062 | 0.173±0.011 | 287.141±14.822 | 0.044±0.001 |
| F | BXD78 | 1 | 13.776 | 10.894 | 1037.972 | 1.974 | 0.162 | 0.164 | 0.087 | 133.266 | 0.038 |
| M | BXD78 | 3 | 14.353±0.252 | 19.580±1.486 | 1158.264±47.949 | 1.691±0.341 | 0.218±0.011 | 0.416±0.056 | 0.138±0.028 | 88.873±39.453 | 0.057±0.010 |
| F | BXD79 | 6 | 12.572±0.534 | 13.090±1.193 | 1076.354±20.095 | 1.361±0.054 | 0.219±0.011 | 0.195±0.009 | 0.078±0.027 | 116.149±54.555 | 0.037±0.003 |
| M | BXD79 | 3 | 13.752±0.145 | 18.117±1.248 | 1086.647± 1.851 | 1.095±0.066 | 0.266±0.006 | 0.384±0.045 | 0.208±0.043 | 161.983± 6.525 | 0.055±0.005 |
| F | BXD83 | 4 | 13.619±0.294 | 14.743±0.985 | 1084.406± 6.020 | 1.492±0.131 | 0.210±0.006 | 0.278±0.020 | 0.103±0.005 | 152.990± 6.569 | 0.041±0.001 |
| M | BXD83 | 6 | 13.880±0.278 | 18.620±1.251 | 1066.919± 6.977 | 1.678±0.156 | 0.225±0.007 | 0.379±0.046 | 0.175±0.024 | 255.159±22.043 | 0.046±0.003 |
| F | BXD84 | 7 | 13.262±0.296 | 12.994±0.753 | 1044.219±13.135 | 1.618±0.087 | 0.194±0.008 | 0.224±0.011 | 0.064±0.017 | 98.203±38.125 | 0.036±0.002 |
| M | BXD84 | 1 | 14.664 | 18.313 | 1005.695 | 1.093 | 0.199 | 0.196 | 0.118 | 187.898 | 0.044 |
| F | BXD85 | 2 | 12.993±0.087 | 13.256±0.306 | 1034.424± 5.131 | 1.628±0.135 | 0.176±0.012 | 0.218±0.041 | 0.096±0.036 | 138.557±29.270 | 0.041±0.004 |
| M | BXD85 | 2 | 13.654±0.085 | 16.545±0.951 | 1054.747± 4.393 | 1.235±0.020 | 0.214±0.004 | 0.303±0.031 | 0.161±0.011 | 184.167±35.782 | 0.048±0.003 |
| F | BXD87 | 2 | 12.079±0.074 | 12.133±0.532 | 1064.210± 6.069 | 1.457±0.080 | 0.205±0.003 | 0.228±0.014 | 0.070±0.001 | 79.452± 4.062 | 0.042±0.001 |
| M | BXD87 | 3 | 12.604±0.293 | 17.495±1.237 | 1067.766±12.969 | 1.183±0.097 | 0.252±0.005 | 0.398±0.005 | 0.166±0.014 | 124.423±15.673 | 0.052±0.001 |
| F | BXD89 | 2 | 12.973±0.018 | 13.266±0.040 | 1062.483± 6.841 | 1.445±0.065 | 0.205 | 0.214±0.001 | 0.064±0.005 | 87.290± 1.713 | 0.037±0.002 |
| M | BXD89 | 6 | 13.092±0.228 | 14.440±0.567 | 1048.943± 6.637 | 1.270±0.041 | 0.230±0.006 | 0.258±0.020 | 0.146±0.013 | 237.926±31.909 | 0.044±0.002 |
| F | BXD90 | 9 | 14.003±0.383 | 17.074±1.174 | 1075.584±12.832 | 1.482±0.100 | 0.201±0.008 | 0.363±0.029 | 0.135±0.015 | 247.976±27.948 | 0.038±0.001 |
| M | BXD90 | 7 | 14.333±0.221 | 18.979±0.524 | 1062.033± 9.645 | 1.432±0.057 | 0.210±0.010 | 0.421±0.034 | 0.170±0.012 | 357.559±27.835 | 0.039±0.001 |
| F | BXD95 | 6 | 13.343±0.299 | 16.776±0.927 | 1037.854±16.495 | 1.466±0.054 | 0.206±0.004 | 0.340±0.025 | 0.131±0.013 | 151.017±16.386 | 0.048±0.002 |
| M | BXD95 | 4 | 13.475±0.305 | 17.151±1.790 | 1022.122±12.965 | 1.517±0.132 | 0.205±0.014 | 0.388±0.051 | 0.144±0.031 | 191.055±41.278 | 0.044±0.003 |
| F | BXD98 | 1 | 14.041 | 16.740 | 1074.541 | 1.211 | 0.193 | 0.264 | 0.148 | 153.834 | 0.051 |
| M | BXD98 | 4 | 12.930±0.380 | 13.346±1.141 | 1048.002±13.947 | 1.557±0.022 | 0.194±0.005 | 0.237±0.013 | 0.093±0.013 | 107.846±19.062 | 0.042±0.001 |
| F | BXD99 | 3 | 12.834±0.208 | 13.892±0.422 | 1027.939± 8.621 | 1.987±0.155 | 0.159±0.003 | 0.188±0.011 | 0.117±0.017 | 234.633±33.424 | 0.038±0.001 |
| M | BXD99 | 0 | N/A | N/A | N/A | N/A | N/A | N/A | N/A | N/A | N/A |
| F | BXD100 | 7 | 13.881±0.283 | 18.234±1.225 | 1035.925±13.065 | 1.318±0.082 | 0.198±0.005 | 0.390±0.024 | 0.150±0.022 | 203.713±28.279 | 0.045±0.001 |
| M | BXD100 | 9 | 14.321±0.197 | 21.895±1.044 | 1068.625±11.070 | 1.666±0.261 | 0.229±0.005 | 0.460±0.029 | 0.194±0.010 | 284.210±14.814 | 0.044±0.001 |
| F | BXD101 | 3 | 13.476±0.394 | 15.290±0.833 | 1046.598± 7.537 | 1.271±0.071 | 0.193±0.005 | 0.226±0.010 | 0.148±0.006 | 158.856± 5.613 | 0.047±0.002 |
| M | BXD101 | 1 | 13.650 | 15.891 | 1026.369 | 1.398±0.111 | 0.184±0.009 | 0.219±0.006 | 0.154±0.017 | 224.839±39.199 | 0.043±0.001 |
| F | BXD102 | 3 | 12.677±0.687 | 14.723±2.329 | 1032.288±23.284 | 1.369±0.074 | 0.188±0.003 | 0.227±0.017 | 0.139±0.029 | 160.763±41.176 | 0.048±0.002 |
| M | BXD102 | 8 | 12.652±0.226 | 14.952±0.697 | 1034.898± 7.545 | 1.396±0.093 | 0.205±0.009 | 0.313±0.014 | 0.130±0.018 | 180.066±20.822 | 0.045±0.002 |
|  | Average<br>per<br>strain | 8.<br>67 | 13.418±0.219 | 15.948±0.856 | 1058.565±10.376 | 1.489±0.114 | 0.206±0.007 | 0.286±0.025 | 0.138±0.017 | 191.835±25.734 | 0.044±0.002 |

Heritability
| Males and Females Combined |  |  |  |  |  |  |  |  |  |  |  |
| --- | --- | --- | --- | --- | --- | --- | --- | --- | --- | --- | --- |
| Variance of Strain Averages |  |  | 0.44 | 5.89 | 833.30 | 0.064 | 4.72E-04 | 0.007 | 0.002 | 5333.20 | 3.19E-05 |
| Total Variance |  |  | 0.69 | 10.14 | 1182.46 | 0.117 | 6.57E-04 | 0.011 | 0.004 | 9087.32 | 4.37E-05 |
| Heritability |  |  | 0.64 | 0.58 | 0.70 | 0.55 | 0.72 | 0.65 | 0.50 | 0.59 | 0.73 |

| <u>Females only</u> |  |  |  |  |  |  |  |  |  |
| --- | --- | --- | --- | --- | --- | --- | --- | --- | --- |
| Variance of Strain Averages | 0.47 | 4.91 | 686.04 | 0.057 | 3.14E-04 | 0.003 | 0.002 | 5132.51 | 1.99E-05 |
| Total Variance | 0.80 | 7.91 | 1311.88 | 0.107 | 5.06E-04 | 0.007 | 0.002 | 7197.09 | 2.98E-05 |
| Heritability_F | 0.59 | 0.62 | 0.52 | 0.53 | 0.62 | 0.46 | 0.69 | 0.71 | 0.67 |
| <u>Males only</u> |  |  |  |  |  |  |  |  |  |
| Variance of Strain Averages | 0.44 | 5.89 | 833.46 | 0.064 | 4.72E-04 | 0.007 | 0.002 | 5333.53 | 3.19E-05 |
| Total Variance | 0.56 | 10.32 | 1139.06 | 0.125 | 6.64E-04 | 0.012 | 0.004 | 8339.11 | 5.32E-05 |
| Heritability_M | 0.78 | 0.56 | 0.73 | 0.52 | 0.71 | 0.61 | 0.48 | 0.64 | 0.60 |

Data are expressed as mean ± S.E. Femur cortical porosity, thickness and polar moment of inertia (pMOI) are representative traits of femoral cortical bone; trabecular bone fraction, connectivity density and thickness are representative traits of trabecular bone.
Broad-sense heritability ( $h^2$ ) = variance of strain averages / total phenotypic variance.

Differences of BXD strain means are more modest than those among all individuals (Figure 3). Typical CV range from a high of 34% for femur trabecular connectivity density to a low of 2% for tibia material BMD. The overall 3-fold difference in bone volume of two extreme cases is reduced to 1.5-fold (13.1 vs 20.2 mm) when the analysis is restricted to strain means. Femoral length ranges from 12.2 ± 0.1 mm in BXD27 to 14.7 ± 0.13 mm in BXD55, representing a ~20% difference. Similar to individual data, there is wider variation in strain means of femoral mineralized volume, with 1.55-fold increase from BXD27 (13.08 ± 0.44 mm3) to BXD55 (18.14 ± 0.74 mm3). Not surprisingly, the variation in BMD is much smaller (~1.7%).

**Figure 3.**
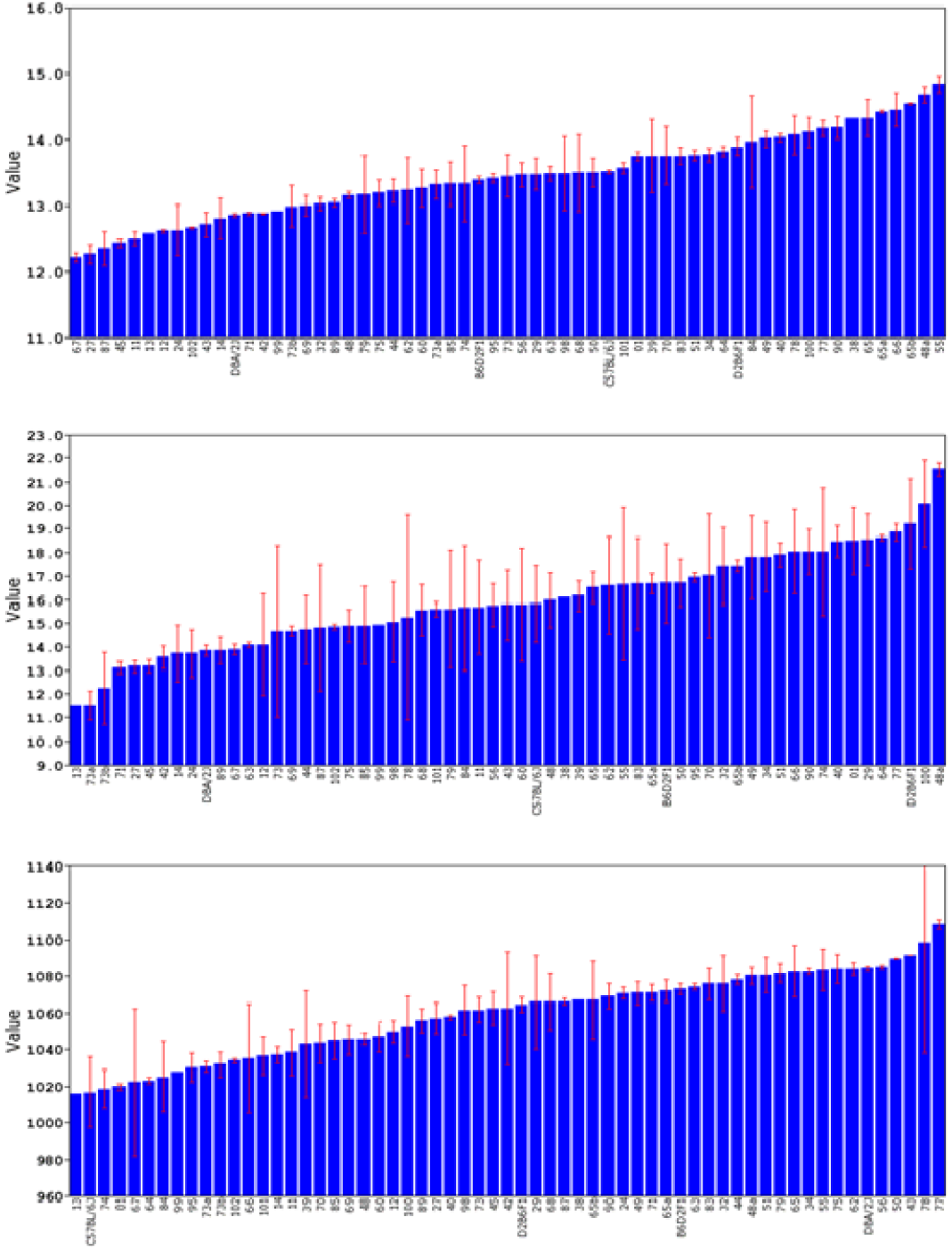
Representative femur traits (sex-averaged) of two parental strains (B6 and D2), reciprocal F1 hybrids (B6D2F1 and D2B6F1) and 59 BXD strains (mean ± SEM). (A) Femoral length (GN 18130). (B) Femoral mineralized volume (GN 18131). (C) Femoral volumetric material BMD (GN 18132). Note that BXD13 is an outlier and requires special handling. We opted to winsorize this value in GN from 963 to 1,016 mgHA/cm^3^.

Femoral length and volume of B6 are greater than those of D2 (Figure 3, 13.5 ± 0.02 mm vs.12.9 ± 0.03 mm in length and 15.8 ± 1.6 mm^3^ vs. 13.8 ± 0.2 mm^3^ in volume, respectively). The material BMD of B6 (1,016 ±19.1 mgHA/cm^3^), however, is lower than that of D2 (1,084 ±1.2 mgHA/cm^3^). A similar pattern is also seen in other strains such as BXD100 and BXD71, but not all strains.

### BXD strain effects and heritability

Among these three key parameters we have studied--strain, sex and age--mouse strain is the dominant factor controlling trait variation. Under carefully controlled laboratory conditions approximately one-third of variation is accounted for by strain even after controlling for log of age and sex. Heritabilities range from 0.29 to 0.78 across all traits (Table 2). The range for females is from 0.36 to 0.69 (mean of 0.59 ± 0.02) and that for males is 0.29 to 0.78 (mean of 0.57 ± 0.01). In our study the typical sample size within strain by sex is 4 to 5. Belknap provides estimate of the corrected or effective heritability of strain means at different resampling depths. When a base heritability of 0.5 is calculated using standard methods then the effective heritability of the means will be close to 0.8 ^(47)^.

Heritabilities were also computed using a jackknife procedure, but did not differ appreciably from those computed using conventional methods (the range is slightly smaller–from 0.25 to 0.67). However, the jackknife procedure also allowed us to estimate standard errors and coefficient of errors of *h^2^* estimates. Given the large sample size (63 strains) and the good control over environmental factors, estimates are surprisingly accurate, and have coefficients of error (SE/mean) that average 1.8%. The highest error is 4.3%, indicating that the heritability estimates are reliable.

### Sex differences

While heritabilities of male and female traits are closely matched (males 0.57 ± 0.01 vs. female 0.59 ± 0.02, mean ± S.E., *r* = 0.29, *n* = 50), values in bone traits differ significantly. With few exceptions, body weight and bone traits are greater in males than females (Table 2). For example, body weight, bone length, and bone volume (Figure 4A) are significantly higher in males. Most other microtraits share this pattern (Figure 4C and 4E). A few traits that do not show sex bias are ratio-based measurement such as femur material BMD (GN 18182 vs. 18232), cortical bone porosity (GN 18185 vs.18235), and trabecular degree of anisotropy (DA, GN 18204 vs. 18254).

**Figure 4.**
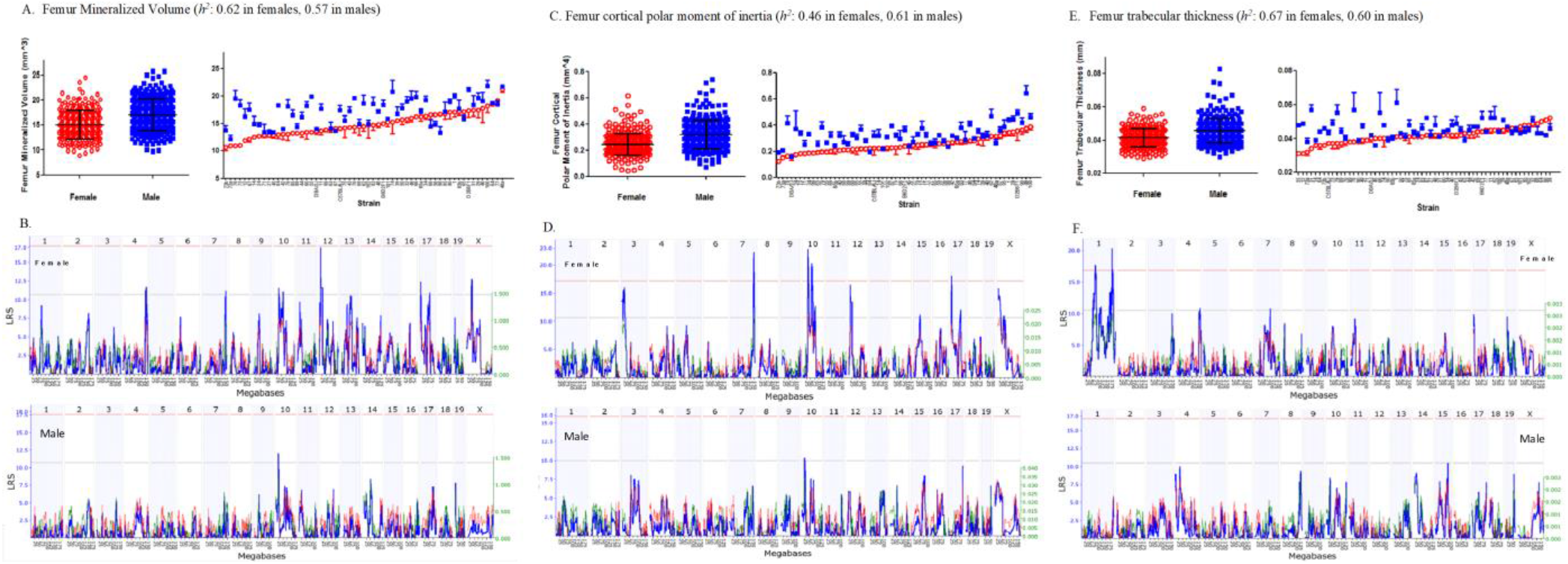
The scatter plots of femur volume (A), cortical bone polar moment of inertia (C) and trabecular thickness (E) of females (red) vs. males (blue). Both the mean values of all animals and most strain averages are higher in males. This pattern is consistent in femur volume, cortical pMOI and trabecular thickness, and vast majority of other microtraits. All the corresponding genomic QTLs (B, D, F) show different locations, peaks and LRS scores between females and males. Data are expressed as mean ± SEM.

### Correlational statistics of bone traits and other phenotypes

Our comparisons are based on nine examples of bone traits—three sex-averaged representative traits each from whole femur and tibia (GN 18130, 18131, 18132), cortical bone (GN 18134, 18136, 18141), and trabecular bone (GN 18146, 18148, 18149). Overall, these sets of whole bone parameters (bone length, bone mineralized volume, and material BMD) correlate modestly with each other (Table 3). However, femur mineralized volume (GN 18131, range from 10 to 21 mm^3^) correlates reasonably well with the other eight traits (Pearson product-moment correlations from 0.28 to 0.70).

**Table 3.** Correlation matrix for 9 key bone traits in femur and tibia

|  | F_Length<br>(GN:18130) | F_Mineralized<br>Volume<br>(GN:18131) | F_BMD<br>(GN:18132) | F_Cortical<br>Bone Volume<br>(GN:18134) | F_Cortical<br>Thickness<br>(GN:18136) | F_Cortical<br>pMOI<br>(GN:18141) | F_Trabecular<br>Bone Fraction<br>(GN:18146) | F_Trabecular<br>Bone SMI<br>(GN:18148) | F_Trabecular<br>Number<br>(GN:18149) |
| --- | --- | --- | --- | --- | --- | --- | --- | --- | --- |
| F_Length | -- | 0.784** | 0.215 | 0.165 | -0.038 | 0.164 | 0.278* | -0.286* | 0.279* |
| F_Mineralized Volume | 0.731** | -- | 0.169 | 0.551** | 0.217 | 0.537** | 0.534** | -0.495** | 0.378** |
| F_BMD | 0.277* | 0.275* | -- | 0.111 | 0.343** | -0.006 | 0.068 | -0.091 | 0.065 |
| F_Cortical Bone Volume | 0.180 | 0.625** | 0.269* | -- | 0.711** | 0.868** | 0.248* | -0.231 | 0.030 |
| F_Cortical Thickness | -0.041 | 0.295** | 0.405** | 0.758** | -- | 0.408** | 0.138 | -0.068 | -0.048 |
| F_Cortical pMOI | 0.184 | 0.575** | 0.115 | 0.899** | 0.474** | -- | 0.234 | -0.239 | 0.071 |
| F_Trabecular Bone Fraction | 0.233 | 0.604** | 0.128 | 0.361* | 0.240 | 0.312* | -- | -0.890** | 0.808** |
| F_Trabecular Bone SMI | -0.254* | -0.549** | -0.166 | -0.282* | -0.151 | -0.262* | -0.899** | -- | -0.672** |
| F_Trabecular Number | 0.219 | 0.418** | 0.029 | 0.117 | 0.006 | 0.124 | 0.824** | -0.652** | -- |

|  | T_Length<br>(GN:18155) | T_Mineralized<br>Volume<br>(GN:18156) | T_BMD<br>(GN:18157) | T_Cortical<br>Bone Volume<br>(GN:18159) | T_Cortical<br>Thickness<br>(GN:18161) | T_Cortical<br>pMOI<br>(GN:18166) | T_Trabecular<br>Bone Fraction<br>(GN:18171) | T_Trabecular<br>Bone SMI<br>(GN:18173) | T_Trabecular<br>Number<br>(GN:18174) |
| --- | --- | --- | --- | --- | --- | --- | --- | --- | --- |
| T_Length | -- | 0.570** | 0.155 | 0.176 | -0.116 | 0.139 | 0.109 | -0.003 | 0.143 |
| T_Mineralized Volume | 0.603** | -- | 0.050 | 0.672** | 0.048 | 0.523** | 0.344** | -0.290 | 0.172 |
| T_BMD | 0.341** | 0.214 | -- | -0.040 | 0.147 | -0.046 | -0.018 | 0.015 | -0.041 |
| T_Cortical Bone Volume | 0.154 | 0.646** | -0.079 | -- | 0.406** | 0.846** | 0.220 | -0.135 | 0.038 |
| T_Cortical Thickness | -0.116 | 0.099 | 0.076 | 0.456** | -- | 0.129 | 0.067 | -0.015 | -0.144 |
| T_Cortical pMOI | 0.138 | 0.498** | -0.038 | 0.783** | 0.111 | -- | 0.137 | 0.006 | 0.063 |
| T_Trabecular Bone Fraction | 0.183 | 0.496** | -0.013 | 0.230 | 0.137 | 0.095 | -- | -0.863** | 0.718** |
| T_Trabecular Bone SMI | -0.117 | -0.446** | -0.112 | -0.150 | -0.011 | 0.010 | -0.852** | -- | -0.515** |
| T_Trabecular Number | 0.163 | 0.319** | -0.035 | 0.033 | -0.042 | -0.009 | 0.829** | -0.584** | -- |

Summary of associations among the most representative bone traits in both femur and tibia. The lower left half lists Pearson product-moment correlations, whereas the upper right half lists the Spearman rank order correlations. All correlation was computed from strain averages.
F: Femur T: Tibia BMD: bone mineral density pMOI: polar moment of inertia SMI: structure model index
\*\* $p < 0.01$ \* $p < 0.05$

Cortical parameters, including cortical volume, thickness, and polar moment of inertia (pMOI), correlate well with each other. The correlation between cortical volume (GN 18134) and pMOI (GN 18141) is 0.90, whereas that between cortical thickness (GN 18136) and pMOI is 0.47. These traits also correlate with whole bone volume and BMD. For example, correlations between total femur volume and cortical volume is 0.63 (GN 18131 and 18134); that between femur BMD and cortical thickness is 0.41 (GN 18132 and 18136), and that between total femur volume and cortical pMOI is 0.58 (GN 18131 and 18141). This is not a surprising finding because cortical bone comprises the largest fraction of mouse long bone. In contrast, trabecular bone parameters—bone fraction BV/TV (GN 18146), SMI (GN 18148), and trabecular number (GN 18149)—do not correlate well with whole bone and cortical bone parameters, except for whole bone volume (range from 0.32 to 0.50). Both the femoral and tibial traits have the same pattern. This site-specificity suggests that the cortical and trabecular components are differentially modulated by gene variants.

We computed correlations between femur traits and other musculoskeletal traits, as well as ~5000 other phenotypes that have already been measured in the BXD family ^(17)^ (**Supplemental Data S2**). As expected, femur length correlates with body weight and length with correlations close to 0.5 (*n* of 20 to 30 common strains). In comparison, the correlation within our own study using precisely the same animals is not much higher (*r* = 0.55 with 397 cases).

MicroCT estimates of femur mineralized volume correlate well with previous DXA data of Jiao and colleagues (DXA, *r* ~0.5, 48 strains in common). However, bones used by Jiao overlap to some degree (about 250 common cases) with those used in the current analysis. Likewise, our femur BMD data correlate to other phenotypes in musculoskeletal system, such as muscle mass ^(48, 49)^ and alkaline phosphatase (ALPL) activity ^(15)^. These bone traits also correlate to several other metabolic traits, including carnitine levels, blood glucose and corticosterone in response to ethanol ^(50, 51)^, body weight gain on high-fat diet, and both high-density lipoprotein (HDL) and low-density lipoprotein (LDL) cholesterol levels ^(52, 53)^.

### Principal component analysis

To systematically analyze the correlation structure among femoral and tibial traits s, we used conventional principal component analysis (PCA) to reduce the dimensionality of 100 traits—50 from males and 50 from females. The first principal component (PC1, GN 18424) accounts for 23% variance and is generally associated with bone volume in both sexes. PC2 and PC3 (GN 18425 and 18426) account for 15% and 12% of variance, and are related to trabecular traits in both femur and tibia, whereas this correlation from PC3 is not strong. PC4 (GN 18427) accounts for 8% of variance and is non-specifically related to the bone length in both femurs and tibias. PC5 and PC6 (GN 18428 and 18429) accounts for 6.5 % and 5% variance, and are generally associated with material BMD in both bones. We also conducted QTL mapping of these top principal components and they did not match up to individual trait well with only one exception: PC4 and femur length in males (GN 18427 and 18230) both map to Chr13 at about 100 Mb. Overall, it was difficult to identify PCs with coherent sets of bone traits, either by sex, or bone type (femur vs. tibia) and site (cortical vs. trabecular).

### Genetic correlations of bone traits with gene expression

We used GeneNetwork to generate lists of the top 1,000 transcripts with expression levels that correlate highly to femur phenotypes using femur mRNA databases (GN410). Lists of these top-ranked transcripts (mean expression level > 7.0) were exported to WebGestalt (WEB-based GEne SeT AnaLysis Toolkit: www.webgestalt.org) for enriched Gene Ontology (GO) category with its sub-root (biological process, molecular function, or cellular component) ^(40, 54)^.

Whereas there were a large number of correlated transcripts, we looked the transcripts with high significant correlations and selected a manageable number to present in **Supplemental Data S2**, giving the priorities to the genes encoding extracellular bone matrix, calcium modulating molecules, receptors, second messengers and relevant hormonal agents and cytokines. While most of related genes are associated with general biological process and molecular function, some are closely relevant to bone biology. For example, the *bone morphogenetic protein 2* (*Bmp2*) and *7* (*Bmp7*) are correlated to femur trabecular fraction (BV/TV), together with *interferon induced transmembrane protein 1* and *5* (*Ifitm1* and *Ifitm5*). And all of these genes are grouped as regulators of ossification and bone mineralization. Additionally, there are other transcripts with high genetic correlation, and encoding protein molecules relevant to bone biology, including TNFRSF, NFκB, insulin, calcium channel, cadherin, and extracellular matrix molecules such as collagen, integrin, fibronectin, hyaluronic acid.

### Genome-wide QTL mapping

We generated QTL maps for all phenotypes (GN 18130 – 18279) and identified a total of 16 significant loci lon Chrs 1, 2, 6, 7, 9, 10, 11, 12, 13, 17 and X (Figure 1C). There was no association between heritabilities of traits and yield of QTLs.

There were only small differences in QTL peak locations (up to 6 Mb) and LRS scores using the two genotype files. The average maximum LRS scores were 13.4 ± 3.2 using the classic file and 13.7 ± 3.3 using the 2017 file. To compare to almost all other data and published QTLs, we have opted to present results using the classic file, but it is a simple matter to update maps using the newer genotype files in GeneNetwork.

We evaluated effects of age differences on mapping results. We compared mapping results from the complete data set of 597 cases (GN 18130 - 18279, with a log age correction) to those from a subset of 484 cases between 65 and 116 days-of-age, without log age correction (GN 18986-19086). The Pearson product-moment correlations between full data with age correction and the trimmed dataset without age correction range from 0.67 to 0.98 (0.93 ± 0.06 SE, *n* = 25 male traits, 25 female traits).

We performed additional analysis that considered genetic markers, mapping algorithm, and composite interval mapping (see *Methods*). We selected seven common and robust QTLs (Figure 1) for key bone microtraits of biological importance ^(8)^. The maximum LRS for all traits is about 22 while the minimum genome-wide threshold (*p* < 0.05) is about 17. These QTLs account for between 25 to 35% of genetic variance in phenotypes (the *r^2^* between the best marker in Table 4 and the strain mean data). Table 4 also provides information on locations, intervals, maximum LRS scores, additive effect sizes, representative bone phenotypes with GN record IDs of these seven QTLs. We also list the *p* values for gene-by-sex interactions. Five QTLs are related to cortical bone traits, whereas two are related to trabecular traits, including *Fttf1a* on Chr 1 and *Ttsf11* on Chr 11.

**Table 4.**
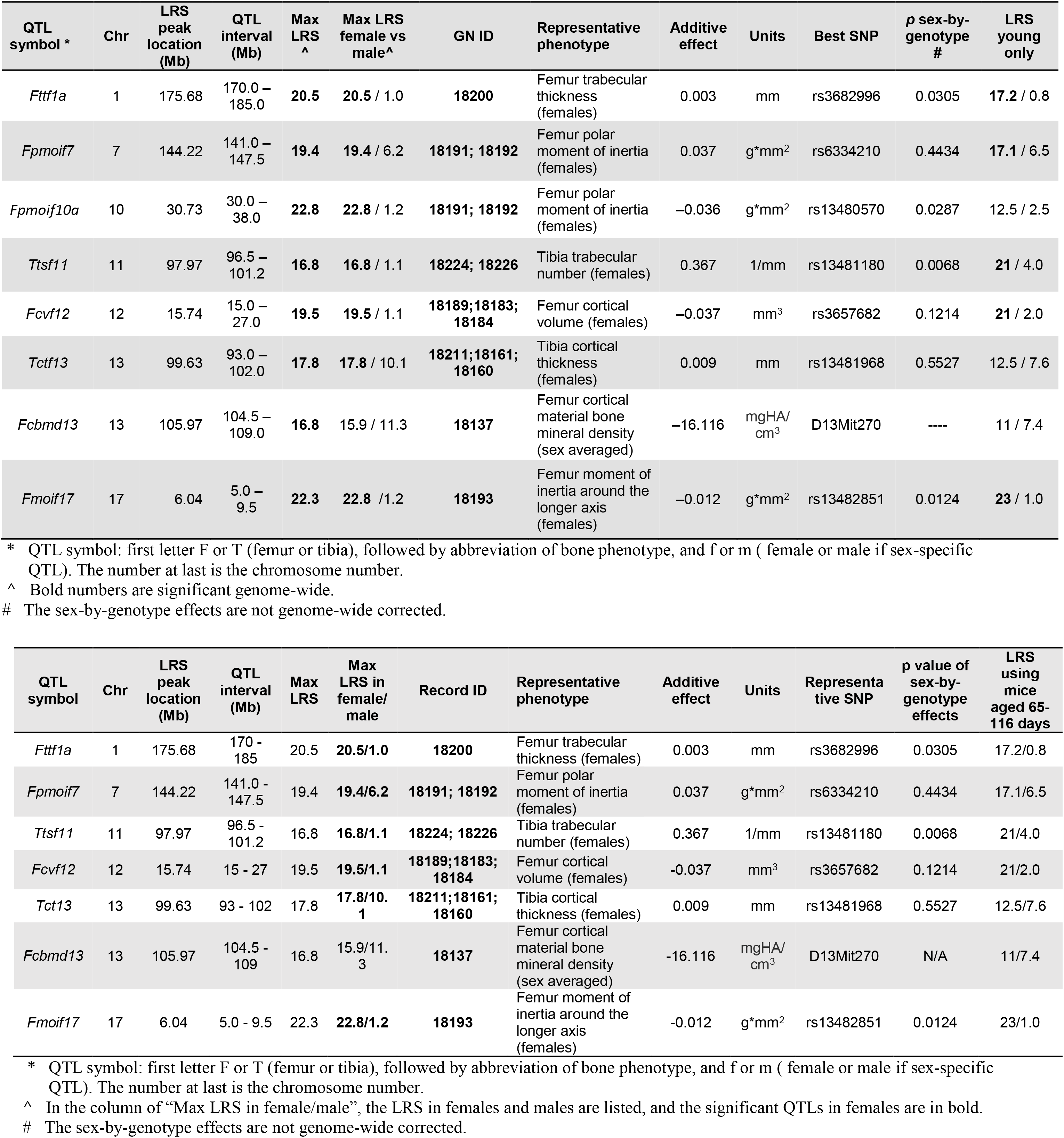
List of 8 strong QTLs, with details of chromosome peak locations, intervals, representative bone traits, sex differences, additive allele effect, corresponding markers and *p* values of gene-by-sex interaction.

#### Sex differences in QTL mapping

We succeeded in mapping significant QTLs for female traits but not for corresponding traits in males. Seven robust QTLs are all detected in females and most of the corresponding QTLs in males are not even suggestive (Table 4). The only exception is tibia cortical thickness in males (GN 18261) reached suggestive level (LRS ~ 11). Among the subset of seven most robust QTLs, four are associated with sex-by-genotype effects.

This sex bias in mapping success was unexpected in our study, because heritability is so closely matched between sexes (**Supplemental Data S1**) though many prior mouse QTL mapping studies have shown a high level of sexual dimorphism ^(55–59)^. We note that body weight loci for the sexes covary well (*r* = 0.69, *n* = 53, GN IDs 18547 and 18548), but QTLs also differ, with suggestive peaks on Chr 1 in males (LRS of 15 at 120 Mb and a high *D* allele) and on Chr 8 in females (LRS of 14 at 80 Mb with a high *D* allele). QTLs for body weight do not match up with any of the bone trait QTLs. And *vice versa*, in comparison with the only male QTL (GN 18248, LRS = 19.7), the LRS of the corresponding female trait (GN 18198) at the same location (~35 Mb on Chr 9) is as low as 0.7. The numbers of the suggestive QTLs in the same interval mappings also share the same pattern, with 42 in females, 19 in males and 31 in sex-averages.

There are several QTLs particularly interesting relevant to sex difference. For example, the femur trabecular thickness in females (GN 18200) has two significant QTLs on Chr 1 (LRS peaks at ~72 and ~176 Mb, respectively), while the corresponding trait in males (GN 18250) is not even suggestive. One candidate gene *Igfbp5* (Insulin-Like Growth Factor-Binding Protein-5, starting at 72.90 Mb, candidate score = 4) in this interval is contributory to skeletal development and BMD acquisition in a sex and age-dependent manner ^(60)^. The distal QTL on Chr 1 also harbors a number of genes modulating bone, including *Grem2, Ddr2, Ifi203, Ifi204, Ifi205* ^(61)^.

There is another strong QTL on Chr 12 for femur cortical volume traits in females only. *Greb1* (growth regulation by estrogen in breast cancer 1) is an estrogen-responsive gene that plays an important role in hormone-responsive tissues and cancer, including breast cancer ^(62)^. It is also expressed in the prostate and its putative promoter contains potential androgen receptor (AR) response elements ^(63)^. Its possible roles relevant to osteocytes and osteoblasts may be mediated by IL6/Stat3 ^(64, 65)^ and Wnt signaling ^(66)^.

Mirroring the results above, the heritability of pairs of male and female traits did not predict whether or not a QTL was detected. QTLs found in females included both cortical and trabecular phenotypes for both bones. Males and females were often littermates, and all phases of the phenotyping were carried out without batch processing by sex. We therefore believe that sex differences in mapping success reflect underlying differences in genetic architecture—for example, traits in males may be controlled by larger numbers of loci with smaller additive effects or controlled to a greater degree by epistatic interactions.

### GO validates known bone-associated genes and defines new bone-associated genes

We extracted sets of genes linked to 34 common bone-associated GO terms from femur gene expression data ^(14)^. These eigengenes correlate well with subsets of known bone-associated genes, but more importantly, also highlight potentially novel genes linked to bone traits. In general, these eigengenes correlate modestly with femur traits and other BXD phenotypes related to the musculoskeletal system. For example, G0:0060346 (bone trabecula formation) includes ten genes: *Chad, Collal, Fbn2, Grem1, Mmp2, Msx2, Ppargclb, Sfrpl, Thbs3, WntlOb*. The eigengenes (GN18479, 18480, 18481) derived from this set correlate with several femoral traits (|*r*| between 0.51 and 0.59), and six tibial trabecular traits (|*r*| between 0.51 and 0.67).

We computed another three eigengenes from G0:0001503 (ossification, see traits GN 18476, 18477, 18478) that also correlate modestly with femoral traits. We selected four known genes for ossification: *Bmp1, Alpl, Mmp23, Colla1* with corresponding transcripts (ILM2940576, ILM2340168, ILM3940278, ILM730020) for correlation comparison analysis to generate 200 probes for this GO. The adjusted *p* values for the top 20 probes range from 10^-6^ (ILM70193 for *Cpz*-carboxypeptidase Z) to 10^-23^ (ILM4780484 for *Pard6g-* par-6 partitioning defective 6 homolog gamma). Among them, 14 genes are known to be implicated to bone biology or encoding bone matrix, including *Bmp1, Alpl, Mmp23, Tmem119, Serpinf1, Ifitm5, Col5a1, Rcn3, Colla1, Adamts2, Smpd3, Pthr1, Mmp16, Col22a1*; and 2 genes (*Maged1* and *Pard6g*) are involved in the regulation of osteoblast activity and/or bone mass ^(67)^. The rest of other genes are not known to be bone-associated, but with a significant GO enrichment *p* value for ossification, including *Fkbp10* (*p<* 10^-22^), *Kdelr3* (*p<* 10^-28^).

We extended this analysis to 34 bone ontology terms, and listed the *p* values and defined –log_10_ (*p*) as the “**bone scores**” (both the average and best score) in each GO in **Supplemental Data S3**. We found there are more than 500 probes with highest *p* value less than 1×10^-17^, i.e., highest bone scores greater than 17. Among them, there are 188 probes with the average bone scores greater than 7.0, and we focused analysis on these genes.

We also counted the total presence numbers or “hits” of each of these 46,621 gene probe (30,890 unique protein-coding genes) in 34 bone GO terms. Not surprisingly, the probes with more “hits” have higher bone scores. The average score is 2.90 for those probes with more than 10 “hits”; 1.73 and 1.54 for those with 5-9 “hits” and 0-4 “hits”, respectively.

However, in this large set of genes and probes, the majority does not have bone GO “hits”. Among the top 20 genes with the highest bone scores (> 8.96), only three (*Cd276, Bmp3*, and *Satb2*) are listed in current bone GO terms. Another five (*Col15a1, Unc5b, Fam78b, Dlx6*, and *Nkd2*) have been implicated in bone biology or skeletal system development, but are not currently part of any bone-related GO term. Surprisingly, the remaining 12 (*P3h4, Unc5b, Srpx, C1orf198, Gxylt2, Clec11a, Sec31a, Bok, Clec11a, Kdelr3, Prss35, Tmtc2*) with these high bone scores have not been known to be functionally associated with bone biology, and are therefore strong candidate genes that are well worth focused analysis and validation.

Collectively, there are only 36 genes associated with bone GO terms among 1,638 positional candidate genes (~2%) within 16 significant QTLs. There genes include: *Adam8, Adamts14, Asf1a, Bcor, Cadm1, Cartpt, Col13a1, Ddr2, Dscaml1, F2r, Gas1, Gata1, Gja1, H3f3a, Hexb, Hey2, Hnrnpu, Id2, Ifi204, Igfbp5, Ihh, Itgb1bp1, Med1, Nodal, P4ha1, Pcgf2, Ptch1, Pvrl1, Rara, Rgn, Sgpl1, Srgn, T, Thra, Wnt6*, and *Zfp640*. The remaining 98% genes are absent in any of current bone GOs.

We have used three gene datasets to define bone ignorome (Figure 2), with a goal to highlight all members of the large set of positional candidates potentially involved in bone biology. The average bone score of 30890 protein-coding genes (blue line in Figure 5) and 1,344 positional candidate genes (yellow line) are close: 0.85 ± 0.01 and 0.88 ± 0.03, as compared to a higher mean of 1.74 ± 0.07 in 770 known bone GO genes (green line). If a stringent threshold of 3 in bone score is chosen, there is a total of 2075 genes (gray area between blue and green lines) above that level. They may be ignorome bone genes. Not surprisingly, the percentages of genes with bone scores above 3 in these categories are 7.3%, 7.2% and 25.2%, respectively (Figure 5). At more stringent levels of 6 and 9, gene numbers above threshold are 437 and 59, respectively. These highly ranked genes with minimal or no known links to bone biology should be carefully evaluated as candidate genes.

**Figure 5.**
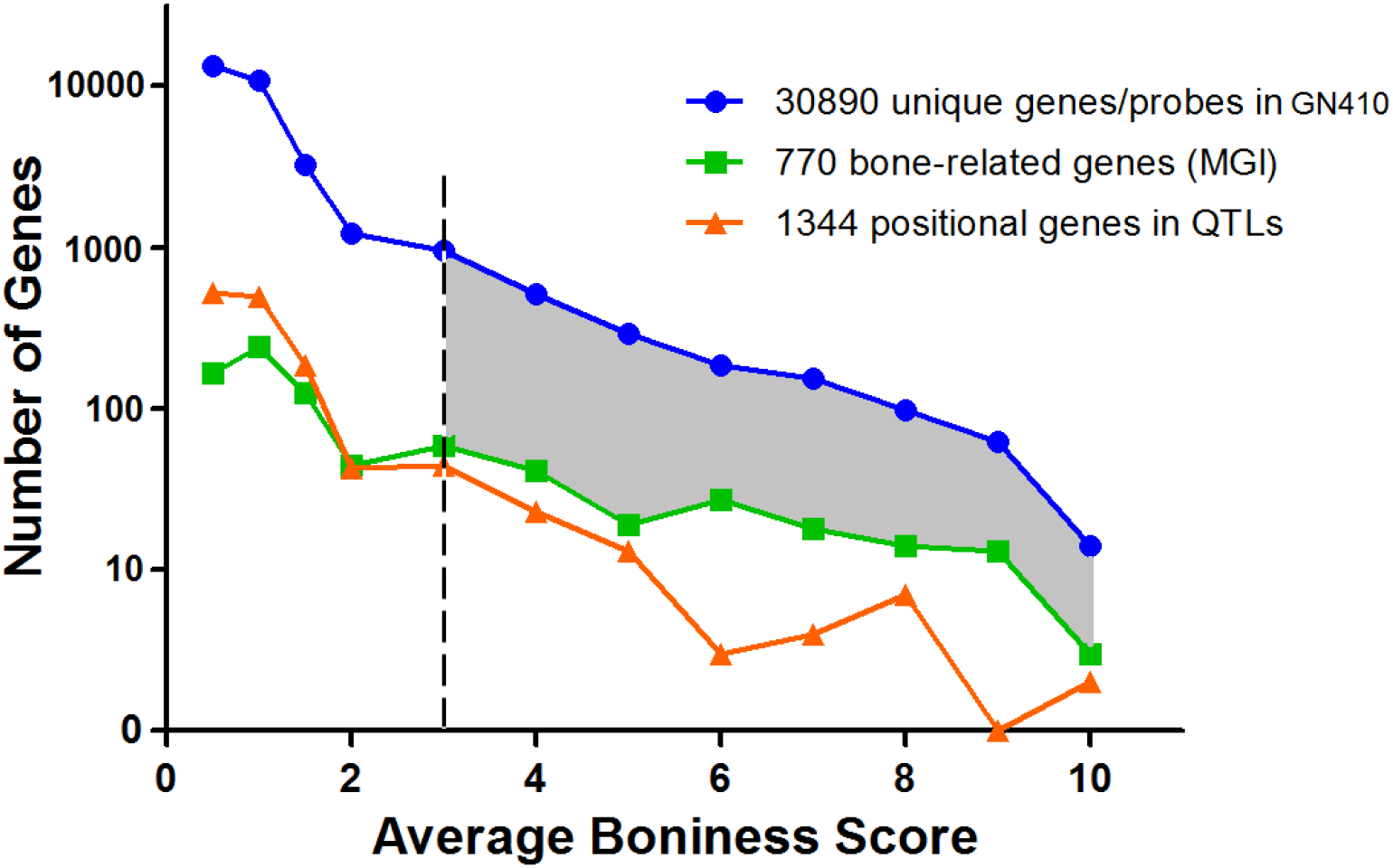
The histogram of average bone scores in three gene sets: 1) blue line: 30,890 unique protein-coding genes in femur mRNA (GN410 dataset with 46,621 Illumina probes); 2) orange line: 1,344 positional genes in 16 QTLs (with bone scores from 1,638 candidate genes); 3) green line: 770 known bone-related genes in MGI. Gray area represents 2,075 protein coding genes without implication with bone biology in current knowledge, but with a bone score greater than 3. Note: the gene numbers on Y-axis is in log10 scale.

#### Candidate genes

We define a total of 212 candidate genes with scores greater than 4. We provide all genome-wide maps, high-resolution maps around peaks, and lists of the top 10 to 20 candidates for each QTL (**Supplemental Data S4**). Among candidates, some are well known genes related to bone biology, such as *Ihh*, interferon activated genes, and genes from the WNT and ADAM families (**Supplemental Data S3**). Some are not well known, but have been linked to rare types of bone disorders. For example, *Fkbp10* (FK506 binding protein 10) is linked to recessive osteogenesis imperfect or Bruck syndrome ^(68–70)^. For the majority of genes, however, their functional linkage to bone biology is largely unknown.

Based on summary candidate scores, we have listed 50 robust candidates with high summary scores between 5 and 8 linked to seven robust QTLs (Table 5). These candidates are grouped into three categories: 1) known bone-associated genes experimentally validated in animal models (*Adamts4, Adam12, Adam17, Ddr2, Darc, Fkbp10, E2f6, Ifi204, Grem2*); 2) candidates with putative bone functions reported in human studies but not yet validated in animal studies (*Ifi202b, Greb1*); 3) bone ignorome genes with high summary scores but no known link to bone (*Ly9, Ifi205, Arhgap30, Slamf9, Ifi203, Sde2, Usp21, Klhdc9, Slamf7, Cd84, Ncstn, Copa, Tmem63a, Ephx1, Cd244, Atp1a4, Slamf8, Pyhin1, Rgs7, Mgmt, Frk, Krt10, Tubg2, Krt12, Stat5a, Trib2, Lpin1, Pqlc3, Hpcal1, F2rl1, Iqgap2, Mrps27, Naip5, Cmya5, Arsb, Polk, Rgnef, Mtap1b, Fndc*)

**Table 5.**
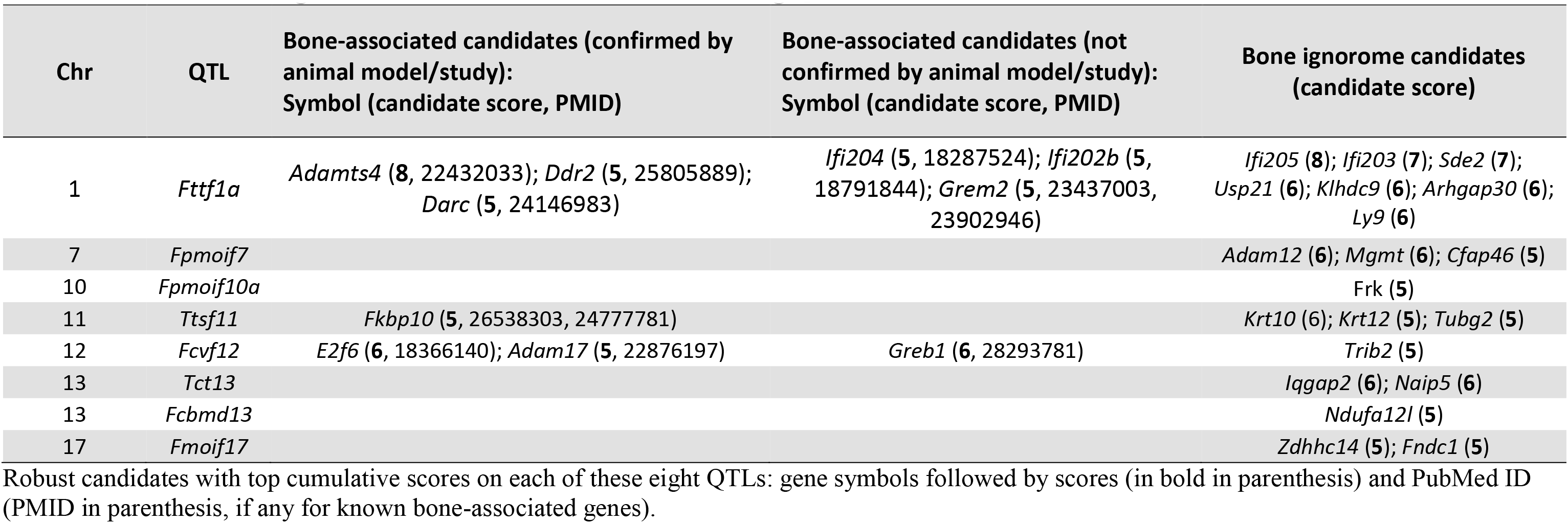
Robust candidate genes based on cumulative scores on 8 strong QTLs.

| Chr | QTL | Bone-associated candidates (confirmed by animal model/study):<br>Symbol (candidate score, PMID) | Bone-associated candidates (not confirmed by animal model/study):<br>Symbol (candidate score, PMID) | Bone ignoreme candidates<br>(candidate score) |
| --- | --- | --- | --- | --- |
| 1 | *Fttf1a* | *Adamts4* (8, 22432033); *Ddr2* (5, 25805889);<br>*Darc* (5, 24146983) | *Ifi204* (5, 18287524); *Ifi202b* (5, 18791844); *Grem2* (5, 23437003, 23902946) | *Ifi205* (8); *Ifi203* (7); *Sde2* (7);<br>*Usp21* (6); *Klhdc9* (6); *Arhgap30* (6);<br>*Ly9* (6) |
| 7 | *Fpmoif7* |  |  | *Adam12* (6); *Mgmt* (6); *Cfap46* (5) |
| 10 | *Fpmoif10a* |  |  | *Frk* (5) |
| 11 | *Ttsf11* | *Fkbp10* (5, 26538303, 24777781) |  | *Krt10* (6); *Krt12* (5); *Tubg2* (5) |
| 12 | *Fcvf12* | *E2f6* (6, 18366140); *Adam17* (5, 22876197) | *Greb1* (6, 28293781) | *Trib2* (5) |
| 13 | *Tct13* |  |  | *Iqgap2* (6); *Naip5* (6) |
| 13 | *Fcbmd13* |  |  | *Ndufa12l* (5) |
| 17 | *Fmoif17* |  |  | *Zdhhc14* (5); *Fndc1* (5) |

Robust candidates with top cumulative scores on each of these eight QTLs: gene symbols followed by scores (in bold in parenthesis) and PubMed ID (PMID in parenthesis, if any for known bone-associated genes).

### Lack of significant epistatic interactions

We searched for two-way epistatic interactions that potentially modulate bone phenotypes in males and female samples. Of a total of 100 traits (50 per sex) we only detected one significant interaction (*p* ~ 0.02) for tibia moment of inertia around the shorter axis in males (GN ID 18267). While this interaction is nominally significant, we are not convinced that this interaction is genuine for two reasons: 1) our search for epistatic interactions does not correct for the many secondary tests we performed, and 2) neither loci in this epistatic pairing are associated with a significant additive effect for this or any of the other 49 traits (Chr 1 at 100.7 Mb and Chr 15 at 33.4 Mb). We therefore conclude that epistatic interactions have relatively modest effects that are not detectible with our modest sample size ^(71, 72)^.

### Combined analysis of mouse candidate genes with QTLs and GWAS hits in rat and human

RGD lists 295 bone QTLs and 646 bone-associated candidate genes in mouse; 213 QTLs and 269 genes in rat; and 70 QTLs and 230 candidate genes in human. In mouse, roughly half of all autosomes are overlapped by one or more of 295 generally broad QTLs.

We compared all 16 of our prime QTLs with those previously mapped in mouse, and found that 14 are novel. Three define entirely new loci (labeled in red in Figure 1C): Chr 6 (*Ttda6*) and Chr X (*FcvfXa* and *FcvfXb*). The others overlap previously reported QTLs but eleven are microCT-derived structural traits not directly related to BMD, and should therefore also be considered novel (labeled in green in Figure 1C), including *Fttf1a, Tcv2, Fpmoip, Ftsmoim9, Ttdaf9, Fpmoif10b, Ttsfll, Fcvf12, Tmoif13, Tct13*, and *Fmoif17*. The only two QTLs with similar phenotypes and map positions, and therefore provisionally replicated, are *Fttf1b* ^(73)^ and *Fpmoif10a* ^(74)^ (labeled in black in Figure 1C).

At the level of candidate genes, only 18 out of 212 of our candidates overlap those already curated in RGD. Of these 18, 17 are genes defined as having a role in bone biology in mouse, 2 in rat, and only 1 (*Ihh*) in human cohorts.

Of 212 candidate genes for all traits seven have human homologs that lie within a GWAS locus as shown in **Supplemental Data S4** (1638_sorted_212candidates_>4): (1) *Grem2* (gremlin 2, DAN family BMP antagonist) (Paternoster et al., 2013) is located on chromosome 1 in both mouse and human and overlaps an association for estimated BMD on chromosome 1 spanning from 240.1 - 241.1 Mbp. *Grem2* is a member of a family of bone morphogenic protein antagonists, is expressed in bone and has been linked to low BMD ^(75, 76)^. (2) *Sorl1* (sortilin-related receptor 1) is a VPS10 multifunctional receptor, serving as a trafficking receptor located in the Golgi apparatus^(77)^. *Sorl1* is located on chromosome 3 in mice, on chromosome 1 in humans, and overlaps an association for estimated BMD on chromosome 1 spanning from 121.3-122.3 Mbp. *Sorl1* has previously been identified in screens for differentially expressed genes in mesenchymal stem cell differentiation, but the mechanism by which *Sorl1* relates to bone biology remains unclear ^(78, 79)^. (3) *Cadm1* (cell adhesion molecule 1) is a cell adhesion molecule that has been observed to be downregulated in many different cancers ^(80)^. *Cadm1* is located on chromosome 9 in mouse, on chromosome 11 in human, and overlaps an association for estimated BMD on chromosome 11 spanning from 115-115.2 Mbp and is just upstream of an association spanning from 115.4-115.6 Mbp. In the context of osteosarcoma, *Cadm1* has been observed on the osteoblast cell surface and has been used as a marker of differentiation in osteoblasts ^(81)^. Additionally, *Cadm1* has been demonstrated to play a role in *NFATc1* regulation of osteoclast activity ^(82)^. However, its role in the determination of bone architecture has not yet been elucidated. (4) *Hap1* (huntingtin-associated protein 1) *Hap1* was identified as interacts with the huntingtin protein and modulates intercellular trafficking in that context ^(83)^. *Hap1* is located on chromosome 11 in mouse, on chromosome 17 in humans, and overlaps an association for BMD on chromosome 17 spanning from 39.4-40.4 Mbp. *Hap1* has not been demonstrated to have a function in bone. (5) *Fancc* (FA complementation group C) is part of a protein complex associated with Fanconi Anemia and plays a role in DNA stability ^(84)^. *Fancc* is located on chromosome 13 in mouse, on chromosome 9 in humans, and overlaps an association for estimate BMD on chromosome 9 spanning from 97.7-98.7 Mbp. *Fancc* has not been demonstrated to have a function in bone. (6) *Ptch1* (patched 1) plays a role in Hedgehog signaling, which is a critical pathway in osteoblast activity (85). *Ptch1* is located on chromosome 13 in mouse, on chromosome 9 in humans, and overlaps an association for estimated BMD on chromosome 9 spanning from 98.1-98.3 Mbp. Mutations in *Ptch1* has been shown to affect bone metabolism ^(86)^ and BMD and fracture ^(87)^. (7) *Msh3* (MutS Homolog 3) is involved in the process of mismatch repair ^(88)^. *Msh3* is located on chromosome 13 in mouse, on chromosome 5 in human, and overlaps an association with estimated BMD on chromosome 5 spanning from 80.2-80.3 Mbp. *Msh3* has not been demonstrated to have a specific function in bone. (8) Tns1 (tensin 1) is located on chromosome 1 in mouse, on chromosome 2 in human and overlaps an association for estimated BMD on chromosome 1 spanning from 218.0 - 218.2 Mbp. Tns1 is concentrated at focal adhesions, binds actin filaments, and regulates cell migration ^(89)^.

In addition, in comparison with the known gene variants with abnormal skeleton morphology on IMPC, there are eight common genes: *Greb1* (score of 6 on Chr 12), *Praf2* (*score of* 4 on Chr *X*)*, Timp1* (*5* on Chr *X*)*, Nr1d1* (*4* on Chr 11), *Thra* (*4* on Chr 11), *Tmem63a* (*5* on Chr 1), *Nsun2* (*4* on Chr 13), *Sdhc* (*4* on Chr 1), with summary candidate scores greater than 4.

## Discussion

### Synopsis

Compared to previous low-resolution studies using DXA, we have used high-resolution microCT imaging to measure bone traits in a large cohort of BXD strains. We selected a subset of 25 traits from trabecular and cortical compartments of tibia and femur for genetic dissection. These microstructural traits have heritabilities that range from 30% to 78% in both sexes. We successfully mapped 16 QTLs—10 for femur and 6 for tibia—and we generated a list of 1,638 positional candidate genes within 1.5 LOD confidence intervals that ranged from 4 to 20 Mb in length. Surprisingly, no QTLs were shared between sexes, and we were far more successful in defining QTLs for females than males, suggesting sex differences in genetic architecture ^(90)^ and the modulation of bone microstructure. From these 16 QTLs we filtered and extracted the seven most consistent traits that we regard to be of highest importance, as well as least dependent on mapping methods or variation in age. We nominated 50 genes with strong associations to skeletal homeostasis and with high summary candidate bone scores using a novel GO annotation strategy. Seven candidate genes have been associated with bone biology or abnormalities in both humans and animal models, including *Adamts4, Ddr2, Darc, Adam12, Fkbp10, E2f6*, and *Adam17*, whereas another four have been linked either in humans (*Grem2* and *Greb1*) or *in vitro* animal models (*Ifi204 and Ifi202b*). All are worth additional genetic and molecular studies to test their roles in bone biology and expression patterns in osteoblasts and osteoclasts. Molecular and cellular functions of the remaining 39 genes are still largely unknown. Some have unusually high bone scores and are therefore primary candidates, especially *Ly9, Ifi205, Mgmt, F2rl1*, and *Iqgap2* that are ranked as top 10 candidates.

The genetic and molecular mechanisms of bone homeostasis, osteoporosis, and other bone diseases are more complicated than originally ^predicted (91, 92)^. However, as we and others have shown, mapping sequence variants modulating bone structure and function is now becoming much easier ^(14, 93–95)^. Defining causal genes is also finally becoming more practical by combining mapping with omics data sets–in our case by combining mapping results with expression data and with GO analyses to generate both bone scores and summary candidate scores ^(96–99)^. The hard job of validation and mechanistic interpretation is also getting easier, with faster gene engineering methods to modify DNA sequence ^(91)^. What is still most challenging is understanding how combinations of loci and gene variants collectively influence bone biology and disease risks and how therapeutic intervention are likely to interact with genotype ^(11, 12, 16)^.

### The significance of computing bone ignorome scores for candidate gene ranking

We compared all 16 of our bone QTLs with 295 published mouse and 213 rat bone QTLs listed in the RGD ^(100, 101)^. Of these 16, three are completely novel—those on Chr 6 (*Tcv2*) and on Chr X (*FcvX* and *FcvfX*). The other 13 overlap bone-associated QTLs reported in RGD, but only two have similar phenotypes and map positions: *Fttf1b* and *Fpmoif10a*. The other 10 of our new QTLs are specifically linked to microCT bone traits and therefore should also be considered novel.

Given the large number of QTLs and positional candidate genes we uncovered, we needed to develop efficient and objective methods to evaluate candidates and their potential role in bone biology. Of 1,638 overlapping the locations of QTLs, only 36 (~ 2%) have already been linked to any of the major bone and skeletal system GO terms (see *Methods*). The major skeletal system GO terms incorporate ~360 of 24,495 genes in the mouse genome (~1.5% of all coding and non-coding genes), and approximately 404 genes when the list is expanded to include rat and human. Similarly, RGD currently lists 652 genes associated with bone structure and function in mouse—roughly 2.7% of all genes. Even this higher value is likely to seriously underestimate the number of genes and the fraction of genome associated with the development, structure, function, and homeostasis of bone.

We therefore needed to develop a more objective and comprehensive way to highlight additional genes potentially associated with bone biology but for which there is currently no compelling experimental or clinical data. In a recent study ^(22)^ we developed a relatively unbiased computational method to generate lists of genes with both high expression and highly selective expression in organs and tissue types using comprehensive transcriptome data. In the present study we have used a new variant of this method to extract a much larger list of genes that have not been previously linked to bone biology. We refer this as the bone ignorome ^(22, 23, 102)^.

The innovation in the present study is that we have defined the bone ignorome by computing a “bone score” based that ranged from a low of 0 to a high of 17. The mean score of known bone genes was 1.74. In comparison, genes without any known association with bone biology in the literature or with bone GO terms had much higher scores. We opted to apply a stringent threshold to define genes with a bone score of 3 or higher as part of a bone ignorome. This scoring system generated a set of 2,075 genes that are potentially associated with bone biology and the skeletal system (Figure 5), a value that we believe is more in line with the numbers of genes likely to have an important and possibly selective effect in bone biology.

### Candidate gene ranking

We have defined strong candidates by combining bone scores above with several other key attributes. This provided us with a relatively objective way to rank candidates ^(16)^. This method can be easily adapted to provide different weighting to different variables, tissues, and crosses.

We reviewed all 16 significant QTLs to evaluate their replicability when using a subset of data with a narrow age range (65 to 116 days) and without age correction (see traits GN 18986-19086). This reduced sample size by a third but did not affect strain number. As expected, mean linkage scores were reduced by the smaller sample size. Seven QTLs are insensitive to age as a confounder, and from these seven robust QTLs, we have nominated 50 candidate genes in Table 5, selecting only those with high summary scores. Two of these are described below as examples of compelling candidates; but this entire set is worth further systematic analysis.

*Grem2* (gremlin 2, DAN family BMP antagonist) that encodes a member of the BMP antagonist family is a strong candidate for femur trabecular thickness at the *Fttf1a* locus on Chr 1. Its effects on BMP signaling and osteoblast differentiation have been confirmed in *in vitro* studies ^(103, 104)^. Genetic variants in the *GREM2* region influence *GREM2* expression in osteoblasts and are associated with fracture risk in humans (3). Another study reported that the minor allele of rs4454537 in *GREM2* is associated with low BMD in hips of a southern Chinese population ^(105)^. Our findings suggest that the BXD family—females in particular—would be good starting point to test genetic and molecular control of *Grem2* and its possible modulation of trabecular thickness.

Another gene, *Greb1*, is a robust candidate for cortical bone volume at the *Fcvf12* locus on Chr 12. This locus also has a strong sex bias with LRS of 19.5 in females but only 1.1 in males. *Greb1* is responsive to estrogen in breast tissue ^(106)^. It is expressed in prostate and its putative promoter contains potential androgen receptor binding sites ^(63, 107)^. In humans, *GREB1* is associated with BMD at two sites with high fracture rates–femoral neck and lumbar spine ^(108)^. However, the association of *Greb1* with bone biology has not been reported previously in animal models. Since both estrogen and androgen are strong regulators in bone remodeling, it is plausible that *Greb1* is a target for both further osteoporotic genetic research and a target for the prevention and treatment of postmenopausal osteoporosis.

In addition to these two obviously strong candidates that already have been characterized in human we also think that the follow genes with both missense mutations and strong cis eQTL would be worth aggressive molecular analysis using CRIPR-Cas9—*Mgmt, Mrps27, Fndc1*, and *Krt10*, that are linked to traits of femur polar moment of inertia, tibia cortical thickness, femur moment of inertia around the longer axis, and tibia trabecular number respectively

### Sex differences

Phenotypic differences of bone traits between sexes are large, but not surprising. Most values for males are greater than those for females, except a few ratio-based traits. However, we were surprised by the marked sex imbalance in numbers of QTLs we were able to maps. While heritability of traits is roughly matched (mean of 0.59 for females and 0.57 for males), the correlations of traits between sexes across all strains is unimpressive—about 0.43 ± 0.02 (mean ± S.E.). The failure to detect QTLs in one or the other sex is not an artifact of the thresholds we used to declare a QTL. For example, femur trabecular thickness in females is linked to two strong and independent QTLs on Chr 1 with LRS values of 16 and 20. In contrast, the same trait in males does not even reach a suggestive level anywhere in the genome, and has a peak LRS of merely 7 on Chr 1. The same sex bias in favor of detectible QTLs in females was also noted among suggestive loci.

We have detected sex-by-genotype interactions for half of these traits, but our analysis is not corrected for a full genome-wide scan. Sex differences in bone are generated by many genetic and non-genetics factors, and detecting interaction effects is difficult even with large sample size. We have reviewed other studies and none have reported such a strong sex by QTL discovery bias ^(109, 110)^.

### Site specificity

Over the past decade more than 150 loci for bone-associated traits have been mapped in many mouse crosses ^(111, 112)^. Causal gene variants have been successfully defined for more than 10 of these, including *Asxl1, Bbx, Cadm1, Cdh11, Fam73B, Prpsap2, Setdb1, Slc38a10, Spns2, Trim45*, and *Trps1* ^(93, 113, 114)^. Most previous work and over 120 of these loci have focused exclusively on BMD, either of the whole body or bone compartments. In comparison to this simple composite measure, we have focused almost exclusively on microCT bone traits from deep phenotyping. But for reference to previous work, we have also included BMD measures. While BMD by DEXA is still the preferred measurement to screen and diagnose osteoporosis, it is a collective “macrotrait” aggregate of both cortical and trabecular compartments, and differs from the fine-grained structural measurements generated by microCT. We measured three microCT-derived BMDs from whole bone, cortex, and trabeculae in ~600 cases, but failed to map any associated QTLs. In contrast, we were able to detect QTLs for several cortical and trabecular traits, particular in females.

Remodeling and turnover of cortical and trabecular (cancellous) bones are differentially controlled and regulated ^(115)^. Trabecular bone is composed of internal rods and plates, forming a lattice that is the primary repository of bone marrow. Because of its close proximity with marrow and marrow-derived cells, trabecular bone has a higher level of turnover than cortical bone ^(116)^. Our genetic dissection of these two compartments confirms the distinction. We performed a correlation test and representative trabecular parameters do not correlate with cortical bone or whole bone parameters (bone fraction BV/TV, SMI and trabecular number). We also conducted PCA, but failed to identify eigentraits that collectively represent these two major bone regions (cortical bone vs. trabecular bone). However, individual trabecular and cortical sites did map well, suggesting that gene variants have relatively precise effects of specific regions and compartments.

We mapped 16 strong QTLs for bone microtraits: six for trabecular bone and ten for cortical bone. There was no overlap among loci. Five of six trabecular bone loci have significant *p* values of sex-by-genotype interactions. In contrast, only two of the femur cortical bone loci have sex-by-genotype effects. Taken together, this confirms that trabecular and cortical bone microtraits are differentially and independently modulated. These findings underscore the importance of precision phenotyping and in mapping and in precision medicine ^(117, 118)^.

### GWAS

Since 2007 over 40 human GWAS studies of BMD have been published on skeletal phenotypes and more than 1000 polymorphisms are associated with BMD ^(1, 6 119–122)^. However, there are no common variants with large effects on BMD or risk of osteoporosis ^(6, 123)^. Minor and major inconsistency are common among individual GWASs and even large-scale meta-analysis. Possible reasons include sample size and power, small effect sizes and population differences, and, variation in statistical procedures ^(124, 125)^. Likewise, almost all of these GWASs are based on areal BMD from DXA measurement at various sites, including lumbar spine, femoral neck, total hip, wrist, radius, and tibia. Paternoster and collegues ^(3)^ published the only GWAS based on peripheral quantitative CT (pQCT) measurement of volumetric BMD (vBMD). The trabecular vBMD was linked to one significant SNP, and cortical vBMD analysis was linked to four loci. This study also confirms that variants related to cortical and trabecular parameters differ—rs1021188 on Chr 13 is associated with cortical porosity whereas rs9287237 on Chr 1 is associated with trabecular bone fraction. Other DXA-based studies ^(126–128)^ have confirmed that BMD variants exert site-specific effects; that is to say that strengths of association and magnitudes of effect differ across different skeletal sites.

### Advantages and limitations

This is one of the first genetic studies of bone microarchitecture in mouse using microCT. The method provides a precise way to quantify and image microarchitecture in trabecular and cortical bone compartments with a resolution of 10 microns or less. In contrast, DXA provides measures of total bone mineral density and bone content. One challenge of microCT is the large number of summary values generated per bone. We chose to evaluate and map 25 traits for each bone that are generally regarded as of genuine significance. This is still a large number, and raises the issue of study-wide false positive rates. All mapping is corrected for genome-wide testing, but is not corrected for numbers of traits–150 total entered into GeneNetwork. As a result a subset of QTLs are likely to be false discoveries. This is the main motivation for why we extracted a core set of seven QTLs that we regard as robust in the sense that they are insensitive to variation in age, genotype file (old versus new), and mapping algorithm (Haley-Knott, R/qtl, pyLMM, GEMMA), or the distribution and transformation of the phenotype (original data or winsorized data to minimize the impact of outliers). There is inevitably still some risk of false discoveries, but we regard these QTLs to be strong enough to warrant independent validation using, for example, CRISPR-*Cas9* engineering, pharmacological manipulation, or in-depth omics analyses. A straightforward alternative at this point would also be to extend the study using an independent panel of BXDs (there are another 80 BXD strains that have not been phenotyped at all), or related genetic crosses such as B6D2 crosses ^(129)^ or Diversity Outbred animals ^(95)^.

Limitations of this study may be obvious. First, we have relatively modest numbers of replicates within strain between sexes. This means that we cannot yet evaluate strain-by-sex effects. However, we are able to evaluate overall sex differences for both phenotypes and QTL maps. Second, the sample size is still too small to detect epistatic interactions.

### Future directions

The fundamental goal of this work is to systematically transition from QTL, to gene, to mechanism, to potential preclinical therapies. The first step is to achieve high quality quantitative measures relevant to bone strength and metabolism and to demonstrate heritable control of variation. The second step is to demonstrate that single loci can be defined with sufficient precision to nominate strong candidate genes. Our work has reached the end of this second stage, but raises questions regarding subsequent steps that can most efficiently validate candidates and test them as therapeutic targets. One approach would produce a multispecies meta-analysis of genes implicated in bone function. Those shared genes across species will be the most relevant candidates. Another consideration is to systematically study gene-by-environmental interactions; something most efficiently handled using cohorts such as the BXDs in which genetically identical cases can be exposed to several environments or treatments. In addition, molecular biological method such as gain- and loss-of-function studies are needed to validate candidate genes and investigate the mechanism. Finally, we need to develop higher throughput ways to test therapeutic interventions starting at early stages and using rigorous quantitative methods; what we should call experimental precision medicine. Again, cohorts of isogenic animals for which we have superb baseline data will be essential resources to achieve this last goal and to evaluate impact of treatment as a function of genotype.

## Acknowledgements

We thank Dr. Aldons J. Lusis for providing gene expression data for GO analysis (UCLA GSE27483 BXD Bone Femur ILM Mouse WG-6 v1.1, GeneNetwork Accession Number: GN410). We thank Lei Yan, Arthur Centeno, and Zachary Sloan for their data entry and technical support of GeneNetwork. This work was supported by Center for Integrative and Translational Genomics (CITG) at the University of Tennessee Health Science Center.

Authors’ roles: Study design: RW, LL, and LDQ. Study conduct: JH, NVD, ZX, FX, WG and LL. Data collection: JH. Data analysis: JH, JW, BZ, OS. Data interpretation: RW, LL, LDQ, CF, and CAB. Drafting manuscript: JH, and RW. Revising manuscript content: CAB, CF, LDQ, LL, and AF. Approving final version of manuscript: LL and RW. RW takes responsibility for the integrity of the data analysis.

## Supplemental Data

Supplemental Data S1: Supplement_Data_S1_BXD_bone_data_Master_Table

Supplemental Data S2: Supplement_Data_S2_Table_correlation

Supplemental Data S3: Supplement_Data_S3_BoneScore_GO

Supplemental Data S4: Supplement_Data_S4_final_candidate_scor

